# Cellular heterogeneity and dynamics of the human uterus in healthy premenopausal women

**DOI:** 10.1101/2024.03.07.583985

**Authors:** Nicole D Ulrich, Alex Vargo, Qianyi Ma, Yu-chi Shen, D. Ford Hannum, Stephen J. Gurczynski, Bethany B. Moore, Samantha Schon, Richard Lieberman, Ariella Shikanov, Erica E. Marsh, Asgerally Fazleabas, Jun Z Li, Saher Sue Hammoud

## Abstract

The human uterus is a complex and dynamic organ whose lining grows, remodels, and regenerates in every menstrual cycle or upon tissue damage. Here we applied single-cell RNA sequencing to profile more the 50,000 uterine cells from both the endometrium and myometrium of 5 healthy premenopausal individuals, and jointly analyzed the data with a previously published dataset from 15 subjects. The resulting normal uterus cell atlas contains more than 167K cells representing the lymphatic endothelium, blood endothelium, stromal, ciliated epithelium, unciliated epithelium, and immune cell populations. Focused analyses within each major cell type and comparisons with subtype labels from prior studies allowed us to document supporting evidence, resolve naming conflicts, and to propose a consensus annotation system of 39 subtypes. We release their gene expression centroids, differentially expressed genes, and mRNA patterns of literature-based markers as a shared community resource. We find many subtypes show dynamic changes over different phases of the cycle and identify multiple potential progenitor cells: compartment-wide progenitors for each major cell type, transitional cells that are upstream of other subtypes, and potential cross-lineage multipotent stromal progenitors that may be capable of replenishing the epithelial, stromal, and endothelial compartments. When compared to the healthy premenopausal samples, a postpartum and a postmenopausal uterus sample revealed substantially altered tissue composition, involving the rise or fall of stromal, endothelial, and immune cells. The cell taxonomy and molecular markers we report here are expected to inform studies of both basic biology of uterine function and its disorders.

**SIGNIFICANCE:** We present single-cell RNA sequencing data from seven individuals (five healthy pre-menopausal women, one post-menopausal woman, and one postpartum) and perform an integrated analysis of this data alongside 15 previously published scRNA-seq datasets. We identified 39 distinct cell subtypes across four major cell types in the uterus. By using RNA velocity analysis and centroid-centroid comparisons we identify multiple computationally predicted progenitor populations for each of the major cell compartments, as well as potential cross-compartment, multi-potent progenitors. While the function and interactions of these cell populations remain to be validated through future experiments, the markers and their "dual characteristics" that we describe will serve as a rich resource to the scientific community. Importantly, we address a significant challenge in the field: reconciling multiple uterine cell taxonomies being proposed. To achieve this, we focused on integrating historical and contemporary knowledge across multiple studies. By providing detailed evidence used for cell classification we lay the groundwork for establishing a stable, consensus cell atlas of the human uterus.

## INTRODUCTION

A distinctive feature of reproductive physiology in some mammals is the menstrual cycle, which occurs in humans, higher-order primates, and certain rodents and bats. It involves shedding of the endometrial lining and its subsequent complete repair, making the uterus one of the most dynamic organs (1). This cyclical regeneration process occurs up to 400 times during a woman’s reproductive lifespan (2–5). Approximately, every 28 days, the cycle initiates with a proliferative phase under estrogen’s influence, marked by a significant thickening of the endometrium from a few millimeters to nearly a centimeter. Following ovulation, the endometrial lining undergoes decidualization in response to progesterone secreted by the corpus luteum in the ovary. This process prepares the endometrium for the potential implantation of an embryo during the secretory phase. In the absence of embryo implantation, the lining sheds back to the basalis layer; and the uterus rapidly undergoes involution, setting the stage for the next cycle (5, 6). This continuous, dynamic remodeling of the uterus is intrinsically linked to systemic hormonal regulation and the ovulation cycle. The cyclic dynamics of the endometrium and myometrium, and their regenerative capacities, raises essential questions about the cellular dynamics during these phases and the identity of the stem or progenitor cells responsible for maintaining uterine tissue homeostasis.

Uterine biology research has increasingly focused on the dynamics of various cell types, including the identify and function of the stem and progenitor cells. Initial studies suspected that these cells reside in the endometrium (7). However, it took several decades before researchers reported the discovery of rare clonogenic epithelial and stromal cell populations in the human endometrium (8, 9). Subsequent studies, either *in vivo* or *in vitro*, have corroborated the presence of stem/progenitor cells that potentially maintain the stroma or epithelium, respectively (4, 5, 10–19). Together, these studies suggested that the myometrium and endometrium compartments are maintained by compartment-specific progenitors.

This understanding becomes more complex due to recent reports of inter-compartment cell lineages. First, in hematopoietic stem-cell (HSC) transplant recipients (XX females receiving XY male HSC) bone marrow-derived human mesenchymal stem cells may contribute to uterine tissue homeostasis (3, 6, 20, 21). Second, it was shown that perivascular or stromal cells in mouse myometrium may undergo a mesenchymal-to-epithelial transition (MET), playing a crucial role in the regeneration of epithelial cells in the uterus of mice after birth (22–24). While molecular markers for these stem/progenitor cells have been identified, our understanding of the cellular diversity in the uterus, especially the dynamics and interdependency of different cell types, remains limited.

RNA sequencing (scRNA-seq) of human endometrium samples has been used to understand uterine tissue heterogeneity (19, 25, 26), with each reporting a cell annotation system that covered diverse stromal, epithelial, and immune cell types. However, a joint analysis is needed to synthesize the multiple taxonomies. Most of the analyzed samples were from endometrial biopsies, thus the knowledge of the basalis endometrium and myometrium is still lacking.

In this study we performed scRNA-seq analyses on full-thickness uterus samples from five premenopausal women, followed by integration with publicly available data for the 15 samples from Garcia-Alonso et al. (25) and Wang et al.(19). This effort led to a consensus cell atlas that includes cell populations present in specific compartments or specific cycle phases. Like past studies, we identified 5 major cell types, comprising lymphatic endothelial, blood endothelial, stromal, epithelial, and immune cell. They were further classified into 39 subtypes, including 8 blood endothelial, 11 stromal, 9 epithelial, and 10 immune cell subtypes. Computational analyses of cellular “velocity” within major cell types revealed putative progenitor cells that may give rise to other cell types. We also present evidence for potential dual-lineage progenitors, such as a myometrial multipotent mesenchymal progenitor capable of contributing to the epithelium and endothelium to ensure continued uterine cyclicity and uterine health.

## RESULTS

### ScRNA-seq of five endometrial and myometrial samples revealed 5 major cell types and their functional subtypes

We performed single-cell RNA-sequencing on full-thickness uterus samples from five healthy donors: three were in the Proliferative phase, and one each in the Mid-Secretory and Late Secretory phase, respectively (**Figure 1A; Supplemental Table 1A**). We separated the endometrium and myometrium layers, and applied tissue-specific dissociation protocols (**Methods**). For one of the donors (D5) the cell counts were low, and we combined the endometrium and myometrium samples, leading to nine scRNA-seq datasets containing a total of >50,000 cells passing quality filtering (**Supplemental Figure 1A**; **Methods**). Clustering analyses for the nine samples, done for each separately, showed similar patterns of cellular heterogeneity, with reproducibly observed in 9-11 clusters (**Supplemental Figure 1B**). This observation confirmed that combining the samples was appropriate, with batch corrections applied by donor (**Methods**), resulting in a joint dataset containing 5 major cell clusters **(Figure 1B)**. By using both statistically generated (**Figure 1C; Supplemental Table 3**) and established literature-based markers (**Figure 1D**) we annotated the five major clusters as lymphatic endothelial (PROX1, NRP2, PDPN, FXYD6, CD36, CAVIN2), blood endothelial (VWF, PECAM1), stromal (MYH11, ACTA2, COL1A1, THY1, PDGFRB, DES), epithelial (consisting of both ciliated epithelial - EPCAM, CAPS, FOXJ1 and non-ciliated epithelial cells – EPCAM, PAX8), and immune (KIT, PTPRC, CD3, RUNX3, S100A9, FOLR2, CD86) cells. The distribution pattern of the five cell types over samples showed that myometrium samples contributed more of the blood endothelial and stromal cells, while the endometrium samples were dominated by the epithelial cells as expected (**Supplemental Figure 1C; Supplemental Table 1B**). Lymphatic endothelial cells and immune cells showed variable contributions from different samples (**Supplemental Figure 1C; Supplemental Table 1B**). Nonetheless, nearly all samples contributed to each of the five major cell types.

**Figure 1.**
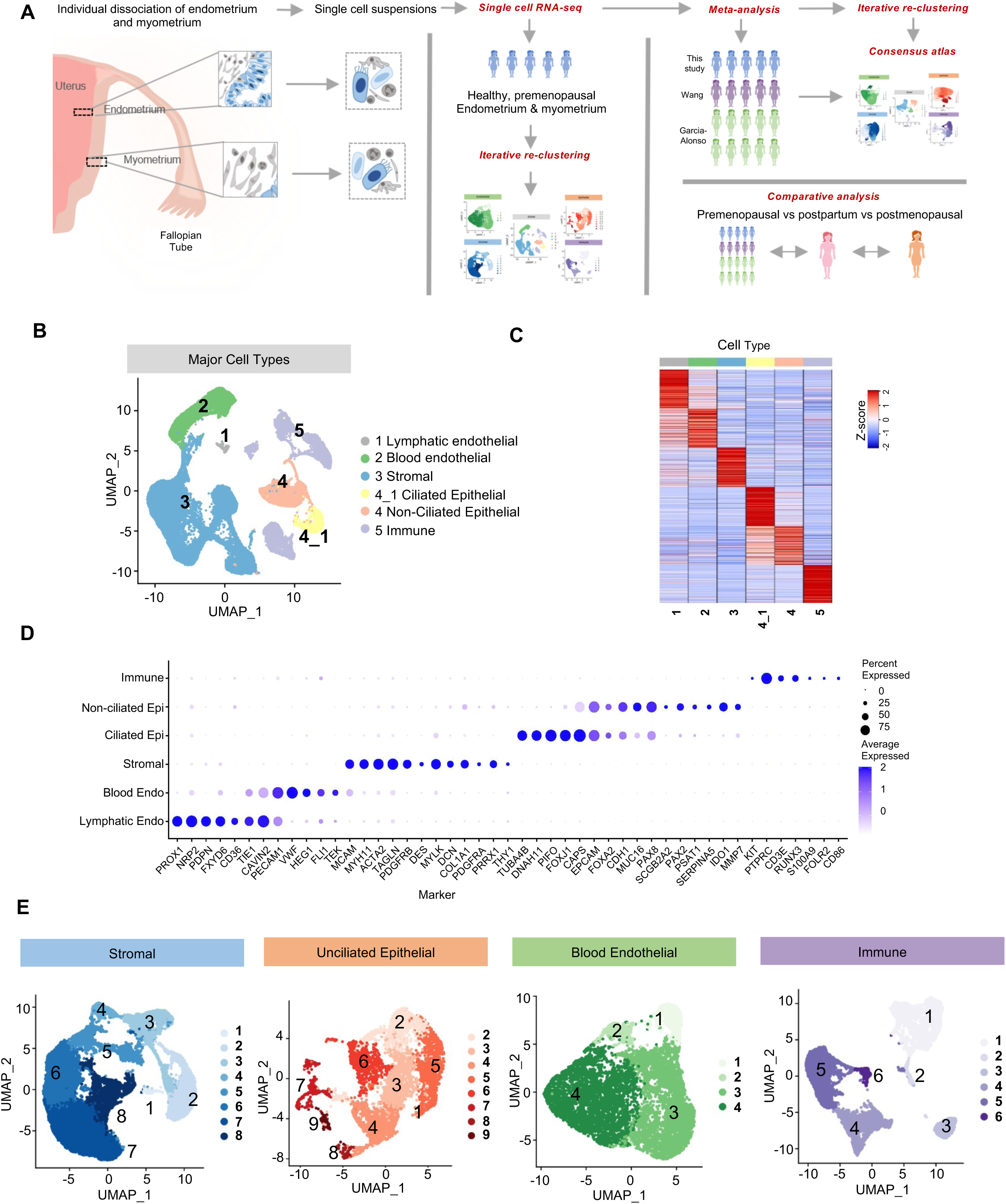
Cell types and markers identified from scRNA-seq analysis of five healthy premenopausal donors. **A**. Overview of the study, including sample and data collection for uterus tissues from the five donors, integrated analyses of prior data from 15 donors to generate a consensus atlas, and comparison with a postpartum and a postmenopausal sample. **B**. Identification of 5 major cell types from global clustering, visualized in UMAP. The epithelial cells are split into ciliated (4_1) and unciliated epithelial cells (4). **C**. Top 150 differentially expressed genes (see Methods) for each major cell types, shown as a gene-centroid heatmap for per-gene standardized values (data in **Supplementary Table 3A**). **D**. Dot plot of literature-based marker genes used to annotate the major cell types. **E**. Identification of cell subtypes by focused re-clustering of each major cell type, shown in UMAP, from left to right, for stromal, unciliated epithelial, blood endothelial, and immune cells.

We subsequently performed iterative re-clustering on each major cell type to discern finer subtypes, except for lymphatic endothelial cells due to the small number of cells recovered. We identified four blood endothelial, eight stromal, nine epithelial (one ciliated and 8 non-ciliated), and six immune cell subtypes (**Figure 1E**), for 27 subtypes (28 if counting lymphatic endothelial). Their cluster centroids and differentially expressed genes, for both the major cell types and the subtypes, were included in **Supplemental Tables 2-3**.

### Joint analyses of 20 samples from 3 studies led to a consensus cell atlas of human uterus

While a natural next step is to biologically annotate the 27 subtypes, we recognized that several prior studies had attempted to define cell types in the human uterus with independent and sometime discordant results. For instance, Wang et al. 2020 (19) generated scRNA-seq data for 10 endometrium samples, while Garcia-Alonso et al. (25) not only collected scRNA-seq data for five new samples in that study (one for both endometrium and myometrium, four with only endometrium), but also attempted a 15-donor joint analysis by combining their 5 donors with the 10 from Wang et al.(19). They reported two endothelial, 7 stromal,10 epithelial, and 14 immune cell subtypes. In a third study, Tan et al. (26) analyzed 14 samples from eutopic endometrial biopsies and ectopic peritoneal or ovarian lesions from patients with and without endometriosis, all undergoing a hormonal treatment, and reported 7 endothelial, 12 stromal, 10 epithelial, and 29 immune cell subtypes.

To reduce confusion among different versions of uterus cell annotation, and to systematically develop a strategy to build a stable, consensus cell atlas, we decided to focus on a multi-study combined clustering and re-clustering analysis. Since the subjects in Tan et al. (26) were either on hormone treatment or diagnosed with endometriosis, we only used their cluster centroids for compariso. For the 15-donor data by Wang et al. (19) and Garcia-Alonso et al. (25) we accessed the individual cell-level data as well as the cell type annotations reported by Garcia-Alonso et al (25). After merging data from the 20 donors (**Supplemental Table 4A**), containing a total of 167,910 cells, we performed data processing and cell filtering using the same standards. After batch-correction by donor, and subsequent clustering, we arrived at five major cell types (**Figure 2A**). They show comparable QC measures (**Supplemental Figure 2A**) and are contributed by all 20 samples (**Supplemental Figure 2B**). Focused re-clustering identified 8 blood endothelial, 11 stromal, 9 epithelial (one ciliated and 8 non-ciliated), and 10 immune cell subtypes (**Figure 2B**), for a total of 38 subtypes (39 if counting lymphatic endothelial). The centroids of major cell types and subtypes were included in **Supplemental Table 5**, with differentially expressed genes shown in **Supplemental Table 6**. **Figure 2C** shows the literature-based marker genes (with underlying data in **Supplemental Table 7**); and **Figure 2D** shows the relative cell proportion over the subtypes, for the stromal, epithelial, blood endothelial, and immune cell compartments separately, and compared across sample groups corresponding to menstrual phases.

**Figure 2.**
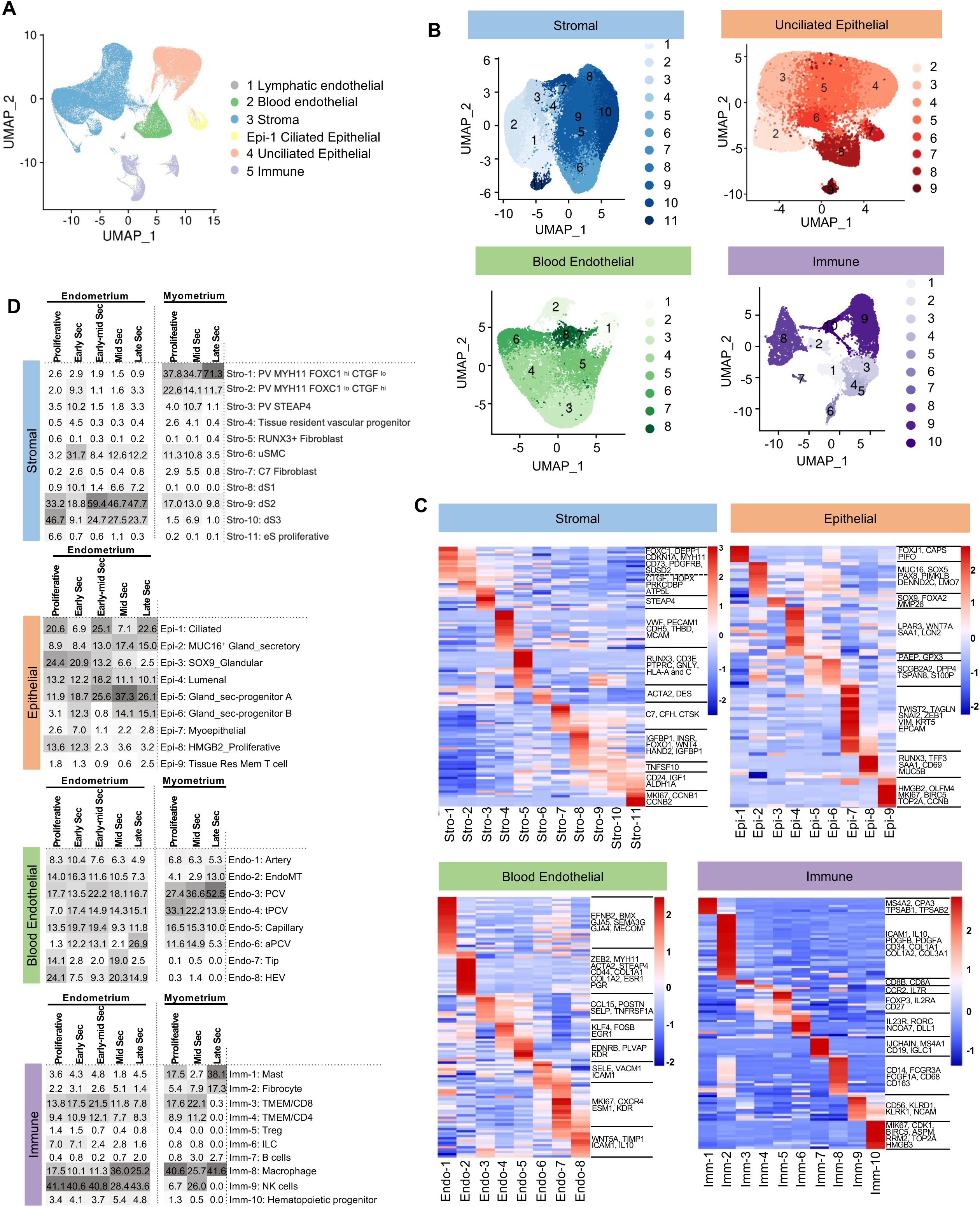
Construction of a consensus cell atlas by joint analyses of scRNAseq data for 20 uterus samples. **A**. Identification of five major cell types, shown in UMAP projection. Epithelial cells are split into the ciliated cells (Epi-1) and the unciliated epithelial cells. **B**. Identification of cell subtypes by focused re-clustering of the four major cell type, shown in separate UMAP plots. **C**. Subtype-specific expression values of literature-based marker genes used to aid in the annotation of the stromal, epithelial, blood endothelial, and immune cell subtypes. Shown are per-gene standardized values from the 20-sample integrated analysis. Data values were in **Supplementary Table 7**. **D**. Comparison of relative abundance of cell subtypes across samples of different cycle phases. Subtype fractions are calculated for the four major cell types separately (top to bottom), and averaged over the samples of the same tissue source (endometrium or myometrium) and the same cycle phase (**Supplemental Table 4**). Values were colored with shades of gray. For epithelial cells only the endometrium samples were shown, as too few cells came from the myometrium samples.

One of the advantages of working with a larger sample size (20 instead of 5) is the improved coverage across the menstrual cycle phases. For example, among our five donors, three were in the Proliferative phase, and one each in the Mid-Secretory and Late Secretory phase, respectively (**Supplemental Table 1A**). In contrast, the phase distribution among 20 donors is much more even representing: Proliferative (7), Early Secretory (4), Early-mid Secretory (3), Mid-Secretory (3), and Late Secretory (3) phases (**Supplemental Table 4A**). Additionally, the joint analyses resolve discrepancies in past cell annotations. For instance, as we show below, two of the stromal subtypes identified in Garcia-Alonso et al.(25), PV MYH11 and PV STEAP4, failed to resemble Prv MYH11 in Tan et al.(26); instead VSMC in Tan et al. (26) strongly resembles PV MYH11 in Garcia-Alonso et al.(25) (see below; **Figure S3B**).

### Challenges in integrated cell annotation and our strategies

The discordant annotations among different cell atlas publications reflected several challenges in the field. First, the biological state of the samples may vary significantly across studies, even with efforts to follow detailed and consistent tissue handling protocols. For example, the Tan et al. (26) samples were collected from subjects on hormonal supplements or with endometriosis. Fourteen of the 15 donor in the joint analysis by Garcia-Alonso et al.(25) only provided endometrium samples. Second, the criteria for cell annotation may rely on a small number of well-known marker genes or on a long list of computationally generated differentially expressed genes, but the relative weights placed on these alternative lines of evidence were not always explained (see the example of SOX9-Proliferative below).

Since the cells from the 15 donors have been clustered by Garcia-Alonso et al. (25) and given names, and they are clustered again in this study in a 20-donor joint analysis, we were able to lay out multiple metrics to examine the level of concordance between the past and new clustering results. In the following sections we present the consensus atlas for the stromal, epithelial, blood endothelial, and immune cells, while applying a consistent set of comparison tools. First, we compare cell cluster assignments between the old and new versions, using *cross-tabulation of cell counts* to indicate the correspondence and stability between two annotation results. Second, we calculate *pairwise centroid-centroid correlation matrices* to measure the correspondence of subtypes between two annotations. Third, we produce panels of differentially expressed (DE) genes from one annotation and plot *gene-by-centroid heatmaps* for centroids from another annotation (and vice versa), to see if the DE genes remain specific in the latter. Since the subjects in Tan et al. (26) were under treatment or having endometriosis, we only applied the second and third approach to the centroids data in Tan et al. (26). With these complementary lines of evidence gathered and compared we were able to propose standardize nomenclature and, importantly, document the ease or difficulty of integrating past results.

### Identification of 11 stromal cell subtypes in the human uterus

The stromal cell compartment makes up the largest proportion of the endometrium and myometrium, and is involved in tissue breakdown and remodeling during the menstrual cycle. Wang et al. (19) analyzed 10 endometrial biopsies and identified a predominant population of stromal smooth muscle cells, which were further classified into two clusters: one consisting of smooth muscle cells that lack the expression of mesenchymal stem cell (MSC) markers; and the other expressing MSC markers such as PDGFRB, MCAM, and SUSD2. Garcia-Alonso et al. integrated these 10 biopsies with an additional five samples and reported seven subtypes: smooth muscle cells (uSMC), fibroblasts C7, two types of perivascular cells (PV STEAP4 and PV MYH11), two types of endometrial cells categorized as decidualized (dS) and non-decidualized (eS) stromal fibroblast, and “other”. Lastly, Tan et al. (26) analyzed 14 diseased or under-treatment samples and reported multiple mural subtypes: vascular smooth muscle cell (VSMC), three perivascular subtypes: PrV-STEAP4, PrV-CCL19, PrV-MYH11, and eight fibroblast subtypes. The latter includes two C7s, two decidualized, and four non-decidualized endometrial fibroblasts.

Focused re-clustering of the ∼89K stromal cells identified 11 subclusters, provisionally labeled as Stro-1 to Stro-11 (**Figure 2B**). Their centroids are in **Supplemental Table 5**. For the stromal cells in our five samples, cross-tabulation of their assignment across the 8 clusters in the initial five-sample analysis and the 11 clusters in the 20-sample joint analysis showed that most of the original 8 clusters were either stably matched or expanded to multiple clusters (**Supplemental Figure 3A**). We then compared the 11 stromal clusters with the clusters reported by Garcia-Alonso et al. (25)and Tan et al. (26), using cross-tabulations (**Supplemental Figure 3D**), cluster-cluster centroid correlations (**Supplemental Figure 3B**), and marker-centroid heatmaps (**Supplemental Figure 3C**). This led to the annotation of Stro-1 to Stro-11 as described below. Their names were shown in **Figure 2D** and **Supplemental Figure 3B**. The expression values for literature-based markers and differentially expressed genes are in **Supplemental Tables 7 and 6**, respectively.

Stro-1 and Stro-2 have high centroid-correlations with PV MYH11 in Garcia-Alonso et al. (25) and with VSMC in Tan et al. (26) (**Supplemental Figure 3B**). Note that while Tan et al. (26) has a PrV-MYH11 population, it does not match Stro-1-2 by either centroid correlation or marker gene expression. Both Stro-1 and Stro-2 express pericyte markers MYH11, PDGFRB, MCAM, RGS5, SUSD2 (**Figure 2C; Supplemental Table 7**). Prior studies have distinguished two varieties of perivascular cells. One has high expression of FOXC1, which promotes pericyte secretion of extracellular matrix (ECM), and could impact the rigidity or mobility of the cells (27, 28). The other has high expression of CTGF, which induces angiogenesis. In our data, Stro-1 expresses higher levels of FOXC1, DEPP1 and CDKN1A, while Stro-2 expresses higher levels of CTGF, HOPX, PRKCDBP (CAVIN3) and ATP5L. We therefore named these two populations as PV MYH11 FOXC1^hi^ CTGF^lo^ and PV MYH11 FOXC1^lo^ CTGF^hi^, respectively. Both are more abundant in myometrium than in endometrium (**Figure 2D**). In endometrium, over the course of the menstrual cycle, Stro-1 peaks in the late secretory phase, while Stro-2 is high in the proliferative phase, suggesting their phase-specific roles during the proliferative and regenerative processes in the uterus.

Stro-3 matches to the STEAP4 perivascular cells in both Garcia-Alonso et al. (25) and Tan et al. (26) for centroid correlations and marker expression (**Supplemental Figure 3B, 3E, 3F**). It has the highest expression of STEAP4 (**Figure 2C**; **Supplemental Table 7**) among all stromal cells and is named <u>PV STEAP4</u>. STEAP4 reduces inflammation and apoptosis in retinal vascular endothelial cells and in many other organs (29). Consistent with a potential immune regulatory role, ligand-receptor analysis indicates that PV STEAP4 cells communicate with macrophages (Imm-8), NK cells (Imm-9) and CD8 cells (Imm-3) (see **Figure 3H** below).

**Figure 3.**
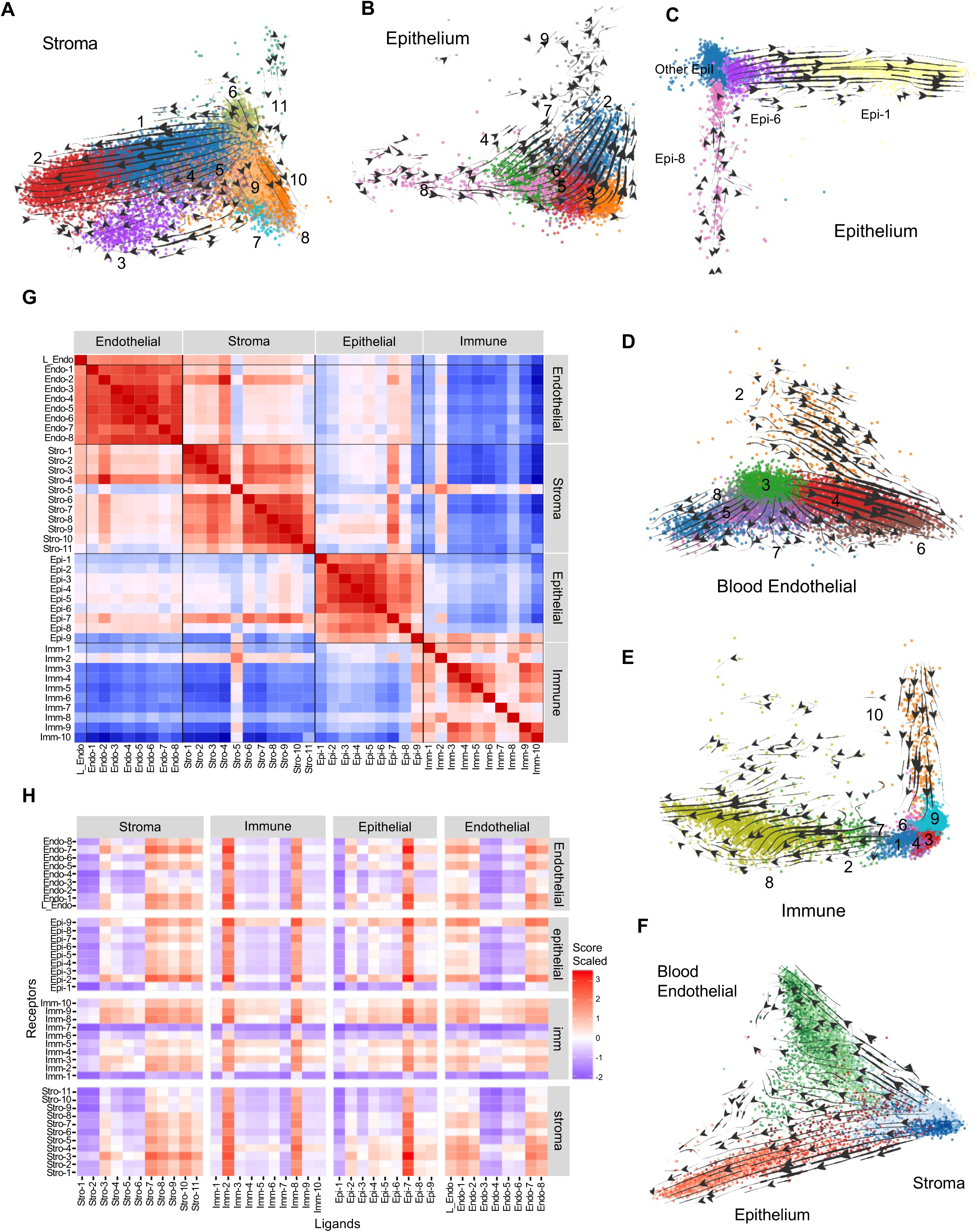
Potential differentiation trajectories inferred from RNA velocity analysis, subtype correlations, and receptor-ligand signaling patterns. **A-F** show velocity analysis (**Methods**) results in PCA projections, with the arrows indicating the local “direction” of transcriptome change. **A**. Velocity patterns among stromal subtypes, showing that Stro-11 is a source population (see main text for more). **B**. Velocity patterns among the unciliated epithelial cells (Epi-2 to Epi-8). **C**. Velocity patterns for epithelial cells after adding the ciliated population (Epi-1, yellow). **D-E**. Velocity patterns among blood endothelial and immune cell subtypes, respectively. **F**. Joint projection of cells in the stormal (blue), blood endothelial (green), and unciliated epithelial (red) compartments, showing that the stromal cells are likely upstream progenitors. **G**. Global comparison of 39 subtypes to highlight dual-character subtypes that may “bridge” across major cell types. Shown in the heatmap of 39-by-39 pairwise rank correlation values among the 39 subtype centroids in the 20-sample analyses. They include the Lymphoid endothelial cells, eight blood endothelial subtypes, 11 stromal subtypes, nine epithelial subtypes (ciliated and unciliated), and the 10 immune subtypes. **H**. Ligand-receptor scores for the 39 subtypes, generally quantifying the communication strengths among them. For each cell type pair we sum the top 5% of potential “interactions” over a curated panel of ∼2000 ligand-receptor pairs (Methods), where “interaction” is the product of logged expression values of the ligand and the receptor in the cell types involved. The subtypes for the ligand is arranged in columns while the subtype for the receptor is in rows.

Stro-4 resembled the smooth muscle cells (uSMC) in Garcia-Alonso et al. (25) in centroid correlation (**Supplemental Figure 3B**) and, to a moderate degree, in cross-tabulation (**Supplemental Figure 3D**). Stro-4 cells express stromal markers in addition to a subset of endothelial marker genes VWF and PECAM1 (**Figure 2C**; **Supplemental Table 7**) but lacks HEG1 and FLI1. Immunostaining confirmed the presence of stromal cells that are VWF+, PECAM1+, but FLI1- (**Supplemental Figure 3H**). Additionally, Stro-4 cells express CD141 (THBD), CD144 (CDH5), CD105 (ENG), MCAM (**Figure 2C**; **Supplemental Table 7**), markers for tissue resident vascular progenitor cells, which are known to reside in the intima of the vessels in individual organs (30). Stro-4 is therefore named <u>tissue resident vascular progenitor</u>. As we show below (see **Figure 3G**), this population appears to bridge with the endothelial cells, especially Endo-MT (see below). Therefore, the identification of potential stromal progenitor of endothelial cells supports a recent model that questions if all endothelial cells come from a bone marrow origin (30, 31).

Stro-5 is a rare cell type and has a poor match between studies, partly due to its small number of cells. It expresses multiple immune markers, including RUNX3 and CD3E, and has similarities with one of the immune cell subtypes, Imm-2: Fibrocytes (see **Figure 3G** below). To verify that the stromal-immune hybrid character of Stro-5 is not due to the population having either stromal or immune cells, we found that individual Stro-5 cells co-express stromal markers (THY1, PDGFRA, COL1A1) and immune markers (RUNX3, PTPRC, GNLY, CD3E) (**Supplemental Figure 3I**). Immunostaining also showed the presence of cells that are RUNX3+, ACTA2+, and PDGFRA+ (**Supplemental Figure 3H**). We therefore labeled Stro-5 as <u>RUNX3+ fibroblast</u>. RUNX3 functions as a tumor suppressor gene in the endometrium and other tissues and is reduced in endometrial cancer. (32, 33). Consistent with a possible role in immune function, Stro-5 expresses the highest level of HLA-A and C (**Figure 2C**; **Supplemental Table 7**), which are class I histocompatibility antigens that present antigen peptides to cytotoxic T cells; we therefore hypothesize that Stro-5 may be a dedicated antigen-presenting fibroblast population.

Stro-6 had high centroid correlation and marker gene similarity with uSMC and “other” in Garcia-Alonso et al. (25) (**Supplemental Figure 3B, 3E**), and is named <u>uSMC</u>. It specifically expresses SMC markers ACTA2 and DES and has moderate expression of MYH11; note that MYH11 is highest in Stro-1/2; **Figure 2C; Supplemental Table 7**). Unlike perivascular smooth muscle cells (Stro-1/2), Stro-5 has much lower expression of MCAM and PDGFRB, two genes highly expressed in PV smooth muscle cells.

Stro-7 matches the centroids and marker expression of Fibroblast C7 in both Garcia-Alonso et al. (25) and Tan et al. (26) (**Supplemental Figure 3B, 3E, 3F**). Notably, C7, the specific marker for this population, was highly expressed in the basal layer of the endometrium as shown by a spatial transcriptomic analysis (25). In our data, Stro-7 is more abundant in myometrium than endometrium samples (**Figure 2D**), possibly due to incomplete separation of the basal layer from the myometrium. Receptor-ligand analysis suggests that C7 fibroblasts communicate broadly, both within stroma and across other compartments (**Figure 3H**),like Stro-8-10, the dS cells.

Stro-8, Stro-9, and Stro-10 matched to decidualized stromal (dS) in Garcia-Alonso et al. (25) (**Supplemental Figure 3B, 3E**) and, somewhat less clearly, to dS1-2 in Tan et al. (26) (**Supplemental Figure 3B, 3F**). The cells express FOXO1, WNT4, and genes involved in insulin signaling (i.e. IRS2, IGFBP1 IGFBP3) (34–36). In addition to cAMP pathway, insulin signaling in response to progesterone has been reported to be a critical pathway for decidualization (35). Insulin signaling activates FOXO1, which is a core transcription factor that regulates downstream decidual marker genes, i.e. IGFBP1, PRL, and WNT4 (**Figure 2C; Supplemental Table 7**; (37, 38)). Given the correlation between datasets and the expression of many decidual cell markers, we label these populations as <u>decidualized fibroblast 1-3</u>. To further distinguish the three subtypes, we noted a prior study reporting that cytokine genes such as CXCL8, CXCL14, and IL6 are part of the senescent associated secretory phenotype (SASP) (39, 40) and are transiently expressed in cells undergoing decidualization. These SASP-related cytokines are expressed higher in Stro-8, while decreasing through Stro-9 and Stro-10. We therefore propose Stro-8 as an <u>early decidualizing</u> state, and Stro-9 and Stro-10 as representing later stages of decidualization.

Stro-11 shows a strong match with endometrial stromal cells (eS) in Garcia-Alonso et al. (25) (**Supplemental Figure 3B, 3E**). The match to the two dS and four eF subtypes in Tan et al. (26) was ambiguous in centroid correlations, except that Stro-11 markers are specifically expressed in eF2 (**Supplemental Figure 3F**). Stro-11 is notable for unique high expressions of cell cycle regulator MKI67 and the cyclins CCNB1-2 (**Figure 2C; Supplemental Table 7**) and is more prevalent in proliferative-stage endometrium samples (**Figure 2D**). We therefore named Stro-11 as <u>eS proliferative</u>. In addition to being the most actively cycling stromal population, velocity analysis identified that Stro-11 is a multipotent progenitor upstream of other stromal cell subtypes (see **Figure 3B** below).

To independently validate the presence of the different stromal subtypes we performed flow cytometry analysis of a full thickness uterus sample using a combination of 7 antibodies (MCAM, CD34, CD24, CD90, CD31, HOPX, CDKN1A). Using the expression patterns of the 8 antibodies we projected thousands of cells using tSNE multidimensional reduction in FlowJo v. 10.10. We then defined 11 groups of cells based on their relative expression levels (table in **Supplemental Figure 3I**) and colored them in the tSNE plot (**Supplemental Figure 3I**), confirming that the existence of the 11 stromal subtypes at the protein level.

In sum, we identified three perivascular smooth muscle cell populations (Stro-1,2,3) that are different from the more abundant uterine smooth muscle cells (Stro-6). Additionally, we identified an unexpected tissue-resident vascular smooth muscle progenitor (Stro-4), which has similarities to endothelial cells (see **Figure 3G** below). Stro-8,9,10 correspond to three stages of decidualized stromal cells; and Stro-11 is a potential proliferative multipotent endometrial stromal cell progenitor (further described in **Figure 3B, 3G**). Stro-5 (RUNX3+ fibroblast) has both stromal and immune cell characteristics; and Stro-7 is a C7 fibroblast that, like other fibroblasts, may recruit various immune cell types (41).

### Identification of nine epithelial cell subtypes in the human uterus

The epithelium layer in the endometrium consists of two major epithelial cell subtypes: glandular and luminal (42). The luminal epithelium cells are involved in embryo implantation, while the glandular epithelium cells produce secretions essential for embryo survival, support decidualization of stromal cells, and regenerate the uterine epithelium after menses (42). To examine the cellular heterogeneity in those two epithelial cell populations, we re-clustered of the ∼47K cells from the 20 donors and identified 9 epithelial sub-clusters, named Epi-1 to Epi-9. To evaluate clustering stability, we compared the cell assignment with the 9 subtypes identified in our original five samples and the new 9 subtypes in the 20-sample analysis. The first two clusters (Epi-1-2) and the last three clusters (Epi-7-9) were relatively similar, while Epi-3-6 had more label shifts (**Supplemental Figure 4A**). These observations were further supported by the centroid-to-centroid correlations between the 5-sample and the 20-sample clustering results (**Supplemental Figure 4B**).

Epi-1 is the most distinct of the epithelial subtypes and readily reproduced over different annotations (**Supplemental Figure 4B-F**). It matches the three ciliated cell types in Garcia-Alonso et al. (25) by cross-correlation, marker expression, and cross-tabulation (**Supplemental Figure 4B, 4C, 4E**) and matches the ciliated population from Tan et al. (26) by cross-correlation and marker expression (**Supplemental Figure 4B, 4F**). It expresses classic ciliated markers FOXJ1, CAPS, and PIFO (**Figure 2C**), and is therefore named <u>Ciliated</u>. Epi-2 had a moderate similarity with Glandular secretory in Garcia-Alonso et al. (25), and with mid secretory in Tan et. al. (26). This population expresses genes such as MUC16, SOX5, PAX8, RIMKLB, DENND2C, and LMO7 (**Figure 2C; Supplemental Table 7**), which are glandular secretory cell markers (43). We labeled it <u>Glandular Secretory</u>, noting that MUC16 is one of the mucins specifically secreted. Epi-3 matched to Glandular and Sox9+LGR5- in Garcia-Alonso et al. (25) and Glandular early-secretary in Tan et al. (26) (**Supplemental Figure 4B, 4E, 4F**). This cluster specifically expresses FOXA2 and MMP26, two known glandular markers (14, 44) as well as SOX9. We therefore named it <u>Sox9 Glandular</u>.

Epi-4 matches Sox9-LGR5+ and Luminal 1 in Garcia-Alonso et al. (25) (**Supplemental Figure 4B, 4C**) and the three Luminal clusters in Tan et al. (26) (**Supplemental Figure 4F**, left panels). It expresses LPAR3, WNT7A (45), SAA1, and LCN2 (46, 47) all implicated in luminal epithelial cell function in the endometrium. Specifically, LPAR3 was shown to be necessary for successful embryo implantation and uterine receptivity in mice (48), and WNT7A is known to be secreted by luminal epithelial cells and may have a role in postmenstrual endometrial repair (45, 49). Epi-4 is therefore named <u>Luminal</u>.

Epi-5 and Epi-6 both matched to Glandular secretory in Garcia-Alonso et al. (25) (**Supplemental Figure 4C,** yellow box in **4E**), and mid secretory in Tan et al. (26) (**Supplemental Figure 4F**). They both highly express PAEP, GPX3 and SCGB2A2, known glandular secretory markers. Between the two, Epi-5 expresses higher levels of S100P, while Epi-6 expresses higher levels of TSPAN8, DPP4, as well as a moderate level of ciliate cell markers: FOXJ1, PIFO, and CAPS (**Figure 2C; Supplemental tables 6**). Velocity analysis suggests that Epi-5 is upstream of Epi-2 Glandular Secretory cells (**Figure 3C**), and Epi-6 is a likely progenitor for Epi-1 Ciliated cells (**Figure 3D**). Across the cycle of endometrium, Epi-2 and -5 peak in the early-mid to late secretory phase (**Figure 2D**). In contrast, Epi-6 and Epi-1 had more variability across the cycle (**Figure 2D**), and more sample-sample variation for those in the same cycle phase (**Supplemental Table 4B**). Taken together, we labeled Epi-5 and Epi-6 as <u>Glandular secretory-progenitor A</u> and <u>B</u>, respectively.

Epi-7 did not have a match in Garcia-Alonso et al (25), but matched to Mesothelial in Tan et al. (26), especially for top markers (**Supplemental Figure 4F**, green box). Tan et al. (26) derived their mesothelial cells from the ectopic peritoneal endometrial lesions and the adjacent areas within the peritoneum. In our study, however, cells were derived from normal uterus without known endometriosis. Epi-7 specifically expresses EMT markers such as TWIST2, TAGLN, SNAI2, ZEB1, VIM, suggesting that it may be a <u>myoepithelial</u> cell population (50). We confirm the presence of this population in the epithelium by co-staining for EPCAM and TAGLN (**Supplemental Figure 4H**). Also notable is that AR is uniquely high in Epi-7, while ESR1 is broadly expressed in epithelial subtypes, PGR is specifically high in Epi-3 and Epi-7,8,9 (**Supplemental Figure 2C**). Interestingly, Epi-7 appears to resemble a population we previously identified in the fallopian tube (51), as shown by pairwise centroid-centroid correlations between the 9 epithelial subtypes in uterus and the 6 epithelial subtypes previously identified in fallopian (**Supplemental Figure 4G**). We therefore labeled Epi-7 as <u>Myoepithelial</u>.

Epi-8 matched well with SOX9 Prolif in Garcia-Alonso et al. (25) in centroid correlation, cross-tabulation, and marker patterns (**Supplemental Figure 4B, 4C, 4E**) and with Glandular in Tan et al. (26) (**Supplemental Figure 4F**). This population expresses specifically high levels of transcription factor HMGB2, OLFM4, and cell cycle genes TOP2A, MKI67, BIRC5, and CCNB. Immunostaining showed the presence of cells that are EPCAM+ and MKI67+ (**Supplemental Figure 4H**). Despite the name Sox9_prolif in Garcia-Alonso et al. (25), Sox9 is only specifically expressed in Epi-3 (Sox9 Glandular) and is at much lower levels in the other eight subtypes, including Epi-8. This low expression of Sox9 is also true for cells in the Sox9_prolif population in Garcia-Alonso et al. (25). We therefore renamed this population as <u>HMGB2_prolif</u>. Velocity analysis (see **Figure 3C, 3D** below) suggests that Epi-8 is the progenitor population for the epithelial compartment.

Epi-9 had no match by centroid correlation (**Supplemental Figure 4B**), but highly expresses RUNX3 and CD69, two tissue resident T-cell markers (**Figure 2C, Supplemental Table 7**). Epi-9 resembled a tissue resident memory T cell population, cluster 2-6, previously identified in the epithelium of the fallopian (**Supplemental Figure 4G**) (51). We confirmed the presence of EPCAM^+^ and RUNX3^+^ cells in the uterus (**Supplemental Figure 4H**). We therefore named this population <u>Tissue resident memory T cell</u>. We show below that Epi-9 appears to be a terminal population in velocity analysis (**Figure 3C**) and has a strong similarity with the immune cells (**Figure 3G**).

Uterine epithelial cells play a crucial role in regenerating the epithelium after shedding and preparing the endometrium for implantation. They accomplish this by communicating with stromal cells to initiate decidualization, thereby preparing the uterine lining for successful implantation (52–54). Mouse models have shown that many pathways, involving estrogen, progesterone, IHH, and BMP, are important for stromal cell decidualization, endometrial receptivity, and embryo implantation (53, 54). Here we find that BMP1 ligands/receptors are lowly expressed in epithelial cells (**Supplemental Table 5**), and ESR1 and PGR are widely expressed across epithelial and stromal cell populations (**Supplemental Figure 2C**). In comparison, IHH signaling appears to be more specific. IHH, the ligand gene, is highly expressed in Epi-3, 8 and 9, while its receptor, PTCH1 is highly expressed in Stro-9, -10, and -11, the dS and eS subtypes (**Supplemental Figure 4J**). Downstream of PTCH1, intracellular decidualization regulators such as FOXO1 and HAND2 and their targets IGFBP1 and IL15 are expressed highly in Stro-8 to 10 (**Figure 2C; Supplemental Table 7**), suggesting that the role of IHH in stromal cell decidualization is likely conserved in humans.

In summary, our re-analysis of epithelial cells recovered five of the previously identified epithelial cell subtypes and renamed Sox9 Prolif to HMGB2 Prolif. Further, we identify three new epithelial cell subtypes: a myoepithelial cell (Epi-7), an epithelial tissue resident T cell population (Epi-9), and a MUC16^+^_Glandular_secretory population (Epi-2).

### Identification of a lymphatic and eight blood endothelial cell subtypes in the human uterus

The cyclical process of endometrial shedding and regeneration constitutes a pivotal aspect of female reproduction in primates. Regenerating a functional endometrial layer post-menses relies on angiogenesis, the formation of new blood vessels (3, 55). Initial clustering of our 5-sample data revealed lymphatic endothelial cell (LEC) and blood endothelial cell as two of the major cell types (**Figure 1B-D**), and they were reproduced in the 20-sample joint analysis (**Figure 2A**). Garcia-Alonso et al. (25) did not report the LEC population in their study; and we found only 27 LECs in their “other” category (**Supplemental Figure 5C**). LECs are clearly matched to the LECs in Tan et al. (26) for both centroid-correlation (**Supplemental Figure 5C**) and the markers (**Supplemental Figure 5D**). We did not cluster LECs further due to having only ∼500 cells in the 20 samples.

While LECs are rare, the blood endothelial cells are much more abundant, with ∼8500 cells in our five samples. The 15 samples in Garcia-Alonso et al. (25) contained another ∼8K endothelial cells, which were classified in that study into two subtypes: ACKR1 and SEMA3G. By re-clustering the ∼16.6K blood endothelial cells from all 20 samples we identified eight subtypes, referred to as Endo-1 to -8 (**Figure 2B**). Comparisons with our original four clusters shows that while the original cluster 2-1 corresponds to the new Endo-1, the other three clusters expanded to 2 or 3 clusters in the new 8-cluster result (**Supplemental Figure 5A, 5C**). Since Garcia-Alonso et al. (25) only reported two subtypes, we focused on comparing Endo-1-8 with the six subtypes in Tan et al. (26), with their names shown in **Figure 2D** and **Supplemental Figure 5C**).

Endo-1 is highly correlated with EC-artery in Tan et al. (26) (**Supplemental Figure 5C-D**) and is named <u>EC-Artery.</u> Consistent with this classification these cells specifically express EFNB2, BMX, GJA5(CX40), GJA4, and MECOM (**Figure 2C**), which are markers for endothelial cells in the artery as opposed to those in the vein. They also express SEMA3G, a marker gene used in Garcia-Alonso et al (25) to define one of the two subtypes of Endothelial cells identified.

Endo-3, 4, and 6 are highly correlated with each other, and with the two endothelial postcapillary venules (PCV) populations (EC-PCVs and EC-aPCVs) in Tan et al. (26) (**Supplemental Figure 5C**). PCV are smaller blood vessels on the venous side of the capillary bed and play a critical role in the exchange of nutrients and waste products between blood and surrounding tissue. They are the preferred sites for lymphocyte extravasation through blood vessel walls to enter the surrounding tissue to phagocytose pathogens, release inflammatory mediators, or perform other immune-related functions (56). In response to inflammatory signals (like TNF-a), EC-PCV transition to activated EC-PCV (EC-aPCVs), causes the venules to become more permeable, meanwhile they synthesize and display adhesions molecules such as SELE (CD62E), ICAM (CD54), and VCAM-1(CD104), which allow for a more efficient recruitment of leukocytes (55, 57). In our data, velocity analysis (see **Figure 3E** below) supports the direction of differentiation from PCV to tPCV, and then aPCV. Furthermore, Endo-3 has the highest correlation with PCV in Tan et al (26), and the expression levels of SELE (CD62E), ICAM(CD54), VCAM1(CD104) are the highest in Endo-6 (**Figure 2C, Supplemental Table 7**). We therefore labeled Endo-3 as <u>EC-PCV</u>, Endo-4 as transitioning EC-PCV (<u>EC-tPCV</u>), and Endo-6 as activated EC-PCV (<u>EC-aPCV</u>). Across the menstrual cycle, PCV (Endo-3) and tPCVs (Endo-4) are detected in both the endometrium and myometrium (**Figure 2D**), while aPCVs (Endo-6) are the most abundant in the late-secretory phase in the endometrium, yet at that phase it is the lowest in the myometrium. These patterns suggest differential roles or regulation of EC-aPCVs in myometrium and endometrium.

It was reported that the transition from PCVs to aPCV may be induced by TNF from neighboring immune cells (57). We found that TNF is highly expressed in several of the immune cell subtypes (especially Imm-4, CD4, see below), whereas the receptor for TNF, TNFRSF1A, is specifically highly expressed in the three PCVs (**Figure 2C**), supporting an important role for immune-to-endothelial cell crosstalk in promoting the PCV to aPCV transition (**Supplemental Figure 5F**).

Endo-8 is moderately matched to HEV in Tan et al. (26) (**Supplemental Figure 5C-D**). These cells express high levels of TIMP1 and OSBPL1A, markers for HEV (**Figure 2C**). Although HEVs were first described in lymphoid tissues, more recent studies identified them in non-lymphoid tissues, including the healthy endometrium (58–60). Like PCVs, HEVs facilitate lymphocyte movement into tissues, playing a crucial role in immune surveillance (56). They are much more abundant in endometrium than myometrium (**Figure 2D**), and enrich in both the proliferative and the mid-late secretary phases. This pattern is the opposite of that of Endo-6, aPCV, which is enriched in the early-to-mid secretory phases.

Endo-5 and Endo-7 correlate with EC-Capillary and EC-Tip, respectively, in Tan et. al. (26) (**Supplemental Figure 5C, 5D**). Endo-5 cells express genes involved in focal adhesion and angiogenesis (CAV1, SPP1, and SOX18). Endo-7 cells express genes involved in vascular genesis including ESM1, CXCR4, and ANGPTL2 (61–63). We therefore labeled them as Capillary and EC-Tip, respectively. In our data, EC-TIP (Endo-7) cells are mainly found in the endometrium, and most abundant in the proliferative phase (**Figure 2D**). These EC-Tip cells play a pivotal role in sprouting and creating new blood vessels in the proliferative phase of the cycle by responding to signals such as vascular endothelial growth factors (VEGF; direct) or ESR stimulation (indirect**) (64)**. Interestingly, while angiogenesis peaks in the proliferative phase (high estrogen and progesterone), (**Figure 2D**) all endothelial cells except for Endo-2, (see below) lacked expression of estrogen receptors ESR1, ESR2 and PGR (**Supplemental Figure 2C**). However, Endo-5 and Endo-7 express high levels of KDR (**Supplemental Table 7**), the VEGF receptor, suggesting that KDR signaling directly promotes angiogenesis while the effects of estrogen signaling are likely indirect.

Finally, Endo-2 is the only subtype that had no clear match in past studies. A unique feature of Endo-2 is its high expression of mesenchymal markers such as CD44, ACTA2, ZEB2, MYH11, STEAP4, and Col1A1/2 (**Figure 2C**), and much decreased expression of classic endothelial markers PECAM1 (CD31), TIE1, and VWF (**Supplemental Table 7**). We recognize it as an endothelial cell population undergoing mesenchymal transition and labeled it <u>EndoMT</u>. It was reported that such cells delaminate and detach from the organized endothelial layer, and migrate into the parenchyma (65). EndoMT is unique among endothelial cell subtypes by expressing both ESR1 and PGR (**Supplemental Figure 2C**), thus it is the only endothelial cell subtype responsive to direct estrogen and progesterone signaling.

In summary, we identified eight blood endothelial cell subtypes. While most correspond to those described in Tan et al. 2022 (26), EndoMT is new. Its developmental origin and function are still unknown.

### Identification of ten immune cell subtypes in the human uterus

Immune regulation is a critical component of the normal menstrual cycle. Immune cells are present in all cellular compartments of the uterus and throughout the cycle, and their presence increases in the endometrium in the late secretory phase (4, 6, 55). This increase contributes to menses and subsequent regeneration, coordinated through communication with other cell types, including tissue stem and/or progenitor cells (4). In this study, the ∼8.7K immune cells in the 5-uterus dataset revealed 6 subclusters (**Figure 1D**). Joint analyses with the immune cells from the 15 samples by Garcia-Alonso et al. (25) identified 10 subtypes: Imm-1 to 10 (**Figure 2B-C**). Crosstabulation with the previous 6 clusters showed that Imm-1, 9, and 10 were relatively stable, while Imm-2-8 expanded three of the original clusters (**Supplemental Figure 6A**). Garcia-Alonso et al. (25) only reported two immune subtypes: Lymphoid and Myeloid, with Myeloid matches to Imm-7-8 and Lymphoid matches to the other eight subtypes (**Supplemental Figure 6B-C**). Tan et al. (26) reported 14 lymphoid and 15 myeloid subtypes. Our annotation, described below, relied more on the comparisons with the fine-scale labels by Tan et al. (26)

Imm-1 had a clear match to mast cells in Tan et al. (26) (**Supplemental Figure 6B, 6F**), and they express classic markers including MS4A2, CPA3, EPPN3, and TPSB2 (**Figure 2C; Supplemental Table 7**).

Thus we labeled Imm-1 as <u>Mast cells</u>. They were more prevalent in myometrium than endometrium and were particularly abundant in the late secretory phase (**Figure 2D**). This is consistent with previous reports of their myometrial enrichment and their role in embryo implantation through hormone induced histamine release (25, 66).

Imm-2 matches to macrophages but to a lesser extent than Imm-8 (**Supplemental Figure 6B**). In addition to resembling macrophages, these cells express stromal markers such as FN1, SPARC, CD34, COL3A1 COL1A1, and COL1A2 (**Figure 2C; Supplemental Table 7)**. We therefore named Imm-2 as <u>fibrocyte</u>, a monocyte-derived cell with characteristics of both macrophages and fibroblasts (67). Fibrocytes have not been previously reported in the uterus, but in other tissues they are involved in early inflammatory responses by expressing ICAM1, and in tissue repair by secreting IL10 (67). Both ICAM1 and IL10 were expressed in Imm-2 (**Figure 2C**), although were even higher in Imm-8 (which we label as macrophage, see below). Consistent with a potential role in endometrial regeneration, these cells express pro-angiogenesis genes PDGFA and PDGFB (**Figure 2C; Supplemental Table 7**). Finally, our analysis of ligand-receptor interactions suggests that both macrophage (Imm-8) and fibrocyte (Imm-2) signal broadly to the stromal, endothelial, and epithelial cells (**Supplemental Figure 3H**). This network of interactions may be involved in the “alternatively activated” macrophage-driven mechanism proposed before for anti-inflammatory tissue repair (4, 67).

Imm-3 and Imm-4 correspond to several cell subtypes around CD8 and CD4 in Tan et. al. (26), and Imm-5 matches to regulatory T cells (Treg) (**Supplemental Figure 6B, 6E**). They express high levels of classic markers: CD3D, CD3G for all three cell subtypes; CD8A, CD8B for Imm-3; CD4 and IL7R for Imm-4, and IL2RA and FOXP3 for Imm-5 (**Figure 2C; Supplemental Table 7**), and as a result were named as <u>CD8</u>, <u>CD4</u>, and <u>Treg</u>, respectively. The relative ratios of different T cells are known to vary during menstrual cycle, as they modulate cell proliferation and estrogen/progesterone signaling in the endometrium (60, 68, 69). It has been shown that the transformation of CD4 into Treg during early pregnancy contributes to immune tolerance - the suppression of immune reactions to the implanting embryo (68). In our data, CD8 and CD4 are maintained stably throughout the cycle, except they are much reduced in the myometrium in the late secretary phase (**Figure 3D**). Treg is much rarer, especially in myometrium. These patterns are consistent with their anticipated recruitment role prior to implantation (69).

Imm-6 has a specific match to ILC in Tan et al. (26) (**Supplemental Figure 6B, 6E**) and is named <u>ILC</u>. They are known to appear in endometrium, but with unknown function (70) In our data, Imm-6 is enriched in the proliferative and early-mid secretory phases in endometrium, and decreases afterwards (**Figure 2D**).

Imm-7 aligns well with B cells and Plasma cells in Tan. et.al. (26), and they express markers for both subtypes, including JCHAIN, CD19, MS4A1 (CD20), CD27, TNFRSF17 (BCMA) (**Figure 2C; Supplemental Table 7**). This cluster contains a relatively small number of cells (∼100) and we did not subcluster it further.

Imm-8 resembles macrophage in Tan et al. (26) (**Supplemental Figure 6B, 6F**), and expresses macrophage markers FOLR2, LYVE1, MRC1, CX3CR1, APOE, VEGFA, SPP1 (**Figure 2C; Supplemental Table 7**), and is named <u>Macrophage</u>. Similarly, Imm-9 matched to several NK subtypes in Tan et al. (26) (**Supplemental Figure 6B, 6E**) and are named as <u>NK Cells</u>. They are a type of innate lymphoid cell acting in immune regulation of embryo implantation by communication with uterine stromal cells, and eventually with the trophoblastic cells of the embryo (71). They have a consistent proportion in the endometrium across the cycle, and a transient increase in the myometrium in the mid-secretary phase (**Figure 2D**).

Imm-10 shows a similar match profile as Imm-9 when compared with Tan et al (26) (**Supplemental Figure 6B**), but unlike Imm-9, it specifically expresses cell proliferation markers and apoptosis inhibitors: MKI67, CDK1, BIRC5, ASPM, RRM2, TOP2A (**Figure 2C**) (72–76). Two of the hematopoietic progenitor markers, CD38 and ITGB1, are specifically expressed in both Imm-9 and Imm-10, although at a higher level in Imm-9, suggesting a close lineage relationship between the two (**Supplemental Table 7**). Importantly, velocity analysis (shown below) highlighted Imm-10 as the progenitor population of the immune cells, upstream of Imm-9 (**Figure 3F**). We therefore named Imm-10 as a <u>hematopoietic progenitor</u> population.

### Velocity analysis reveals progenitors in epithelium, endothelium, and stromal compartments

Past studies have shown that the uterus harbors progenitor cells in different tissue compartments, including an epithelial, endometrial mesenchymal stem cells (eMSCs), side population cells, and bone marrow stem cells (12, 20, 21, 77–83). However, it has been difficult to study cell dynamics *in vivo*, as it requires complex experiments such as lineage tracing or imaging of cell type-specific markers. Towards this goal, we sought to understand the potential lineage relationship among uterine cell subtypes by performing RNA “velocity” analysis, which leverages the spliced and unspliced transcripts from RNA sequencing data to infer the cells’ future states. By estimating the direction of change among cells, velocity analysis provides an upstream-downstream ordering of some cell types, potentially reflecting their progenitor-descendant status during differentiation (84).

First, we focused on the major cell types separately, aiming to infer the progenitor population within the stromal, epithelial, endothelium, and immune compartments. Among the 11 stromal subtypes (**Figure 3A**), Stro-11 (eS proliferative) is apparently the “source” - a multipotent progenitor - for other stromal cell subtypes. One path of differentiation goes from Stro-11 to Stro-1 to Stro-2, the two PV MYH11 populations, while others go to Stro-3 (STEAP4) and Stro8-9-10 (the dS subtypes). Stro-4 (tissue resident vascular progenitor) resides in the middle of the paths. Among the eight non-ciliated epithelial subtypes (**Figure 3B**), Epi-8 (Sox9_Prolif) is the likely progenitor, with Epi-5 (Glandular secretory-progenitors A) upstream of Epi-2 (Glandular secretory), which is a terminally differentiated population. Curiously, Epi-9, the Tissue resident T cell, is an outlier, suggesting that its cell of origin may not be from an existing epithelial cell subtype, and we show below that it has a strong immune characteristics. When we added Epi-1, Ciliated cells, to the rest of the epithelial cells (**Figure 3C**), the Ciliated cells descended from Epi-6, Glandular secretory B, and Epi-8 remains the upstream epithelial progenitor. For the eight endothelial subtypes (**Figure 3D**), Endo-3 (PCV) is the source population, which generates the transitioning PCV (Endo-4, tPCV) and subsequently the activated PCV (Endo-6, aPCV). Endo1 (Artery) and Endo-7 (Tip) are the other terminal populations. Endo-2 (EndoMT) appears as an outlier, consistent with its endothelial-stromal dual characteristics. Finally, for the 10 immune subtypes (**Figure 3E**), Imm-10 (hematopoietic progenitor) is the source population, feeding into Imm-9 (NK cells), while Imm-8 (macrophage) is terminally differentiated.

Next, to gauge the global relationship among major cell types, we projected the stromal, epithelial, and endothelial cells together, and observed a strong “flow” where stromal cells are upstream of both epithelial and endothelial cells (**Figure 3F**). This suggests that, unlike mice, human uterus stromal cells have the potential to undergo a mesenchymal-to-epithelial transition to contribute to the epithelial and endothelial cell compartments during the normal menstrual cycle (22–24).

To further understand the potential transitions, we examined the pairwise similarity among all 39 cell subtypes, focusing on identifying populations displaying intermediate properties between major cell types. First, Endo-2 (EndoMT) has similarity to stromal cells, especially Stro-4 (tissue resident vascular progenitor), which in turn has similarities with most endothelial cells. Second, Epi-7 (myoepithelial) has resemblance to epithelial and stromal cells, as well as with Endo-MT cells (Endo 2). These dual-character subtypes may act as cross-lineage progenitors and, as the global velocity analysis shows (**Figure 3F**), likely involve the stromal cells (such as Stro-11) as the upstream progenitors.

In summary, velocity analysis revealed compartment-specific progenitors. The stromal progenitor, Stro-11, may act as a uterus-wide multi-lineage progenitor. Additionally, we identified transitioning progenitors such as Stro-4, Endo-2, and Epi-7, that have dual characteristics of major cell types. These predicted progenitor populations may be involved in complex and coordinated tissue regeneration programs, involving a hierarchy of multiple progenitors for optimal tissue maintenance and regeneration.

### Population shifts in postpartum and postmenopausal states

The uterus exhibits remarkable regenerative abilities. It rapidly recovers its size and function after menstruation and childbirth. After menopause, even when the ovarian reserve is depleted, the uterus can regain function if stimulated by exogenous estrogen and progesterone, enabling it to support embryo implantation and pregnancy (85). The mechanisms governing tissue homeostasis and regeneration during a typical menstrual cycle are not fully understood, nor are the patterns in postmenopausal woman or following childbirth. While building a normal uterus cell atlas we had the opportunity to analyze a rare postpartum uterus sample (Donor-7, one-week postdelivery) and the myometrium from a postmenopausal uterus (Donor-6, age 52), which were used to focus on the composition of the tissue among the cell types.

At the level of major cell types **(Figure 4A**), the postpartum sample showed dramatically expanded immune cells and much reduced stromal cells, consistent with the immune cell involvement in tissue repair as the uterus returns to its normal state after a pregnancy. The reverse trend is seen in the postmenopausal sample: much expanded stromal cells and significantly reduced immune and endothelial cells, indicating a more stable tissue state without the cyclic turnover or rapid regeneration. Both showed very few epithelial cells. Furthermore, the lymphatic endothelial population is significantly expanded in the postpartum sample, as would be expected in its state of tissue repair.

**Figure 4.**
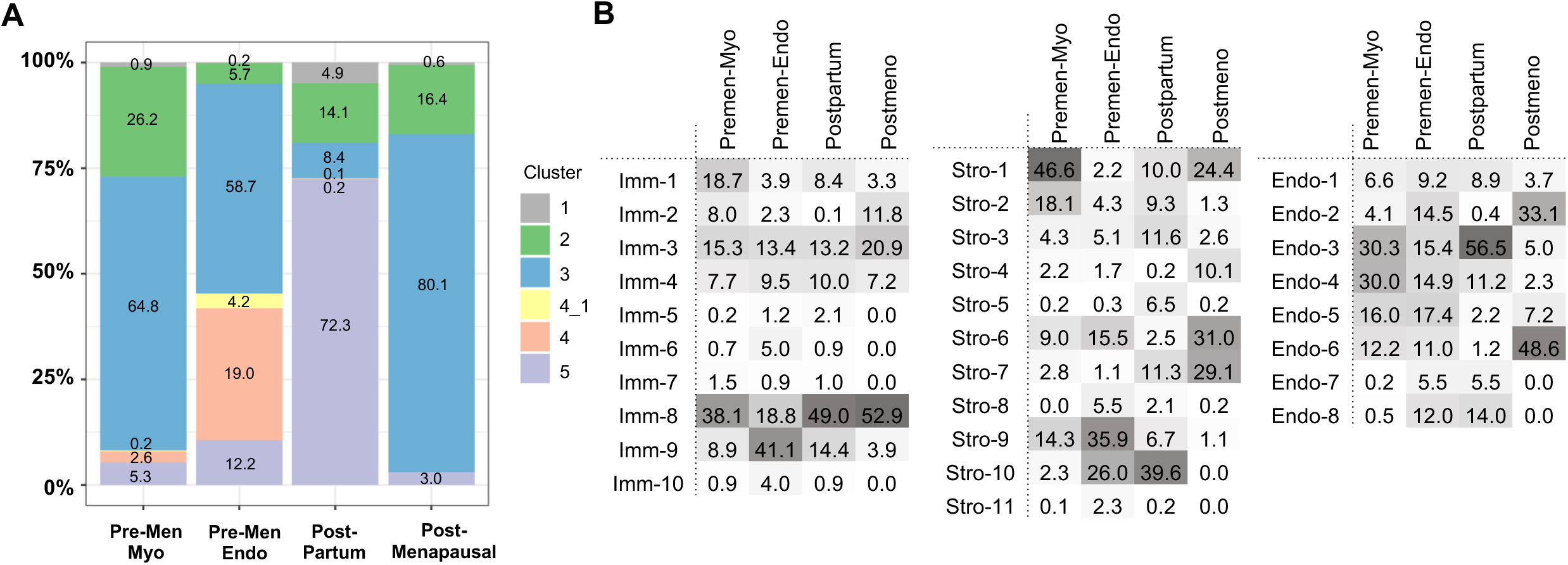
Altered tissue composition by cell types and subtypes in postpartum and postmenopausal uterus samples. **A**. Comparison of relative fractions of the major cell types across four types of uterus samples: endometrium and myometrium samples from the 20 premenopausal donors, a postpartum sample, and a postmenopausal sample. B. Comparison of relative fractions of subtypes, for immune, stromal, and blood endothelial subtypes. Data for individual samples from the premenopausal donors (before being average) are in Supplemental Table 4B.

We then evaluated the relative proportions of cell subtype within each compartment (**Figure 4B**). In the postmenopausal sample. Stro-4 (vascular progenitor), Endo-2 (Endo MT), and Endo-6 (aPCVs) are much more abundant, indicating subtype-specific alterations in vascular composition in postmenopausal uterus. In addition, the Stro-1:Stro-2 ratio (i.e., between the two types of MYH11 subtypes, for ECM secretion and angiogenesis, respectively) is increased, possibly contributing to uterine tissue fibrosis observed in the aging uterus. Finally, it has been shown that during uterine aging, the ability of endometrial stromal cells to decidualize decreases gradually (86) and here we find that the early dS cells (Stro-8) are significantly reduced, while later dS cells, Stro-10, are lost in the postmenopausal sample.

During the postpartum period, the uterus undergoes involution, by which it returns to its pre-pregnancy size. This process involves apoptosis, proliferation, and autophagy of terminally differentiated cells, which are then regenerated as the uterus transitions back to normal cycling (6). In our data, the postpartum uterus exhibits not only a lower proportion of stromal cells (**Figure 4A**), but specifically much lower abundance of Stro-6, the uterine smooth muscle cells. Interestingly, Stro-5 (RUNX3+ fibroblast) is expanded in the postpartum uterus, Since RUNX3 expression in fibroblasts inhibits proliferation, the expansion of Stro-5 may act to prevent the over-proliferation of other stromal subtypes following pregnancy.

In both the postpartum and postmenopausal uterus, Stro-7 (C7 fibroblasts) is increased. Other notable changes are the reduction of Imm-2 (fibrocytes) in the postpartum sample and the reduction of Imm-10 (hematopoietic progenitors) in the postmenopausal sample. The patterns need to be replicated in additional samples, and their functional impacts require further validation.

## Discussion

Investigating the cellular composition and dynamics of the human uterus has been challenging, partly due to cyclic changes in the functional states of many cells, their interactions, and the turnover of cell communities. To profile only the endometrium or the myometrium or having limited samples that fail to cover the menstrual cycle more completely, has hindered a comprehensive understanding of uterus physiology).

In this study, we collected full thickness uterus scRNA-seq data from five donors and integrated with previously published data from 15 donors (14/15 had endometrium only), along with centroid data from a third study. Since each of the prior reports produced a cell annotation system, we sought to examine the concordance and inconsistencies among them, and to uncover potential progenitor populations essential for uterus tissue maintenance and regeneration. For instance, among the epithelial cells, we discovered myoepithelial cells (Epi-7), tissue-resident T cells (Epi-9), which bear resemblance to similar populations (**Supplemental Figure 4G**) found in the fallopian tube epithelium (51), underscoring a shared cellular composition across reproductive tract tissues. Similarly, among the stromal cells, we identified a tissue-resident vascular progenitor (Stro-4), a proliferative mesenchymal progenitor (Stro-11), and RUNX3+ fibroblasts (Stro-5). Furthermore, with the large number of cells from myometrium we were able to further classify the PV MYH11 into two subtypes (Stro-1 and Stro-2), which appear to serve distinct roles in vascular genesis (FOXC1^lo^ CTGF^hi^) and extracellular matrix maintenance (FOXC1^hi^ CTGF^lo^). Among the immune cells we identify a hematopoietic progenitor population (Imm-10) and a fibrocyte population (Imm-2), which has immune-stromal dual characteristics. Lastly, among the endothelial cells, we identified EndoMT cells, which are cells undergoing endothelial-to-mesenchymal transition, and three populations forming a trajectory from basal, transitioning, to activated post-capillary venules (PVC, aPCVC, and tPCV). Taken together, these results led to an expanded consensus cell atlas for the human uterus.

Homeostatic mechanisms in the uterus vary across species and seem to depend on the extent of tissue repair needed (2, 4, 12, 87). For example, in mice, uterine lining regeneration during the normal cycle is maintained by compartment-specific progenitors for the epithelial and stromal cells (22, 88). However, studies using genetic lineage tracing in mice have shown that, after parturition, a subset of stromal cells undergo a MET transition to generate epithelial cells, which are then stably integrated into the luminal and glandular epithelium (22, 88). The molecular identity of the cells undergoing MET was recently shown to be a PDGFRA+ fibroblast population (23). Therefore, tissue homeostasis mechanisms in mouse uterus differ between normal cycles and the extensive regeneration after parturition. In contrast, in vivo lineage tracing analyses are not feasible in humans. Therefore, characterization of stem/progenitor cells has relied on in vitro cell culture assays which has led to the identification of an endometrial mesenchymal stem cells (eMSCs), side population (SP) cells, and bone marrow stem cells with stem/progenitor-like properties(2, 11, 12, 17, 20, 21, 77–82, 89). The precise molecular identities of these cells, how they compare across studies, and their contributions to uterine tissue homeostasis and regeneration remain difficult to define. In this regard we expect that the cellular atlas described here, including both endometrial and myometrial cell subtypes and their mRNA markers, represents a new resource for more detailed explorations into the uterine stem/progenitor cells.

Here, our joint analysis of ∼167K single cells potentially identified multiple distinct compartment-specific progenitors including: Stro-11 (eS proliferative) for stromal cells, Epi-8 (HMGB2-proliferative) for epithelial cells, Endo-3 (PCV) for blood endothelial cells, and Imm-10 (hematopoietic progenitors) for immune cells. Additionally, we uncover unexpected compartment differentiation hierarchy, such as Epi-6 as a likely precursor of the ciliated cells, and Epi-5 (Glandular secretory-progenitors A) as upstream progenitor for Epi-2 (Glandular secretory).

In addition to compartment-specific progenitors, we find that the uterine tissue homeostasis may also rely on cross-compartment, multipotent progenitors. For example, in **Figure 4G** we noted several dual-character populations: Endo-2 (EndoMT), an endothelial cell with high correlations with multiple stromal cell populations, especially with Stro-4 which is a tissue resident vascular progenitor. Similarly, Epi-7 a myoepithelial cell has high centroid correlations to stromal subtypes. Given the identification of these dual state cells, we hypothesized that they may act as cross-compartment progenitors stemming from the stromal compartment; however, their lineage relationships and functional roles remain to be characterized in the future. Importantly, the markers we generated in this study are expected to enable more precise experiments for tracking, isolating, and perturbing these interesting populations. For instance, we find that EndoMT cells are unique among the blood endothelial subtypes in the expression of ESR1 and PGR (**Supplemental Figure 2C**), a feature especially relevant for promoting angiogenesis in the proliferative phase, which is hormonally stimulated. However, the role and significance of this population during vascular remodeling phase is unknown. Similarly, AR is broadly expressed in all endothelial cells and, in other compartments, but appears to be limited to Epi-7 (myoepithelial). AR signaling plays a significant role in endometrial regulation and repair during menstruation; this raises questions about whether Epi7 may have a role in this process. Furthermore, high AR expression in the uterus is also associated with anti-proliferative effects; and its loss in endometrial cancer cells correlates with poorer prognosis (90, 91), raising questions about whether mis-regulation in Epi-7 may be a root cause. In short, these patterns may guide future validation studies in uterine organoid models and allow a more targeted analysis for certain cell types in different disease contexts.

Finally, the recruitment of bone marrow-derived stem cells (BMDSCs) for uterine regeneration was previously reported in female patients receiving bone marrow transplant, where the donor’s bone marrow cells were detected in the endometrium of the recipient (21). In mice, BMDSCs have been shown to contribute to uterine repair (92). In our analysis, while we identified Imm-10, a hematopoietic progenitor population, velocity analysis only suggested its source status among the immune cells, while evidence is lacking for its potential contribution to other compartments. This may be related to its compartment-specific role during normal cycle, while bone marrow cell contribution may only be seen in large-scale regeneration processes after delivery or radiation treatment.

We attempted to use the normal cell atlas to explore tissue-level changes in a postpartum sample and a postmenopausal sample. By examining the differences in cell proportions, we provided an early glimpse of altered tissue function in these atypical states. The observations reported here will need to be replicated in more samples.

Taken together, the cell atlas we describe here provides more than just consensus cell subtypes and markers, but also serves a new foundation for new hypotheses regarding multiple hierarchies of progenitor cells and differentiation trajectories. The detailed transcriptomic profiles of various cell types can serve as a knowledge resource for refining in vitro differentiation protocols and enhance our ability to measure the interactions and dynamics of cell communities in both healthy and diseased states. Being able to understand the lineage and signaling relationships both between and within uterine cell compartments is directly relevant for addressing clinical challenges such as uterine factor infertility, embryo implantation failure, and recurrent pregnancy loss. For example, one of the difficult tasks has been to identify a transcriptomic signature from endometrial biopsies that can predict the window of implantation, the ideal time at which the endometrium is most receptive to embryo implantation, thus improving IVF outcome for patients with implantation failure and RPL (93). The difficulty in identifying such signatures may stem from variations in cell composition within tissue biopsies, which can differ significantly across individuals and in various pathological states. Future research aimed at better understanding the natural heterogeneity of cell subtypes is essential for characterizing an ideal cell state signature. Once established, these signatures have the potential to reveal tissue functionality and ultimately contribute to the development of personalized medicine and targeted treatment strategies. For example, the ratio of senescent decidualized cells to non-decidualized cells in endometrial biopsies has been correlated with RPL (40, 94), highlighting the importance of cellular composition and state in reproductive health. This cell atlas thus stands as a critical knowledge resource, facilitating the advancement of reproductive medicine by enabling a deeper understanding of the lineage and signaling relationships within and between uterine cell compartments.

## Author Contributions

S.S.H. and J.Z.L oversaw project design and analysis. J.Z.L. oversaw computational analysis. Q.M., A.V., D.F.H., and J.Z.L. analyzed data. N.U., Y.S., S.B., S.S.H performed experiments. N.U., S.S.H., and J.Z.L wrote the manuscript with help from coauthors. Comments from all authors were provided.

## Acknowledgements

We thank members of the Hammoud and Li Labs for scientific discussions and manuscript comments. This research was supported by National Institute of Health (NIH) 1DP2HD091949-01 (S.S.H.), T32-GM70449 (D.F.H.), HL144481 (BBM), Open Philanthropy grant 2019-199327 (5384) (S.S.H.) and Chan Zuckerberg Foundation Grant CZF2019-002428 (A.S., E.E.M., J.Z.L. and S.S.H.).

## Supplemental Figure Legends

**Supplementary Figure 1.**
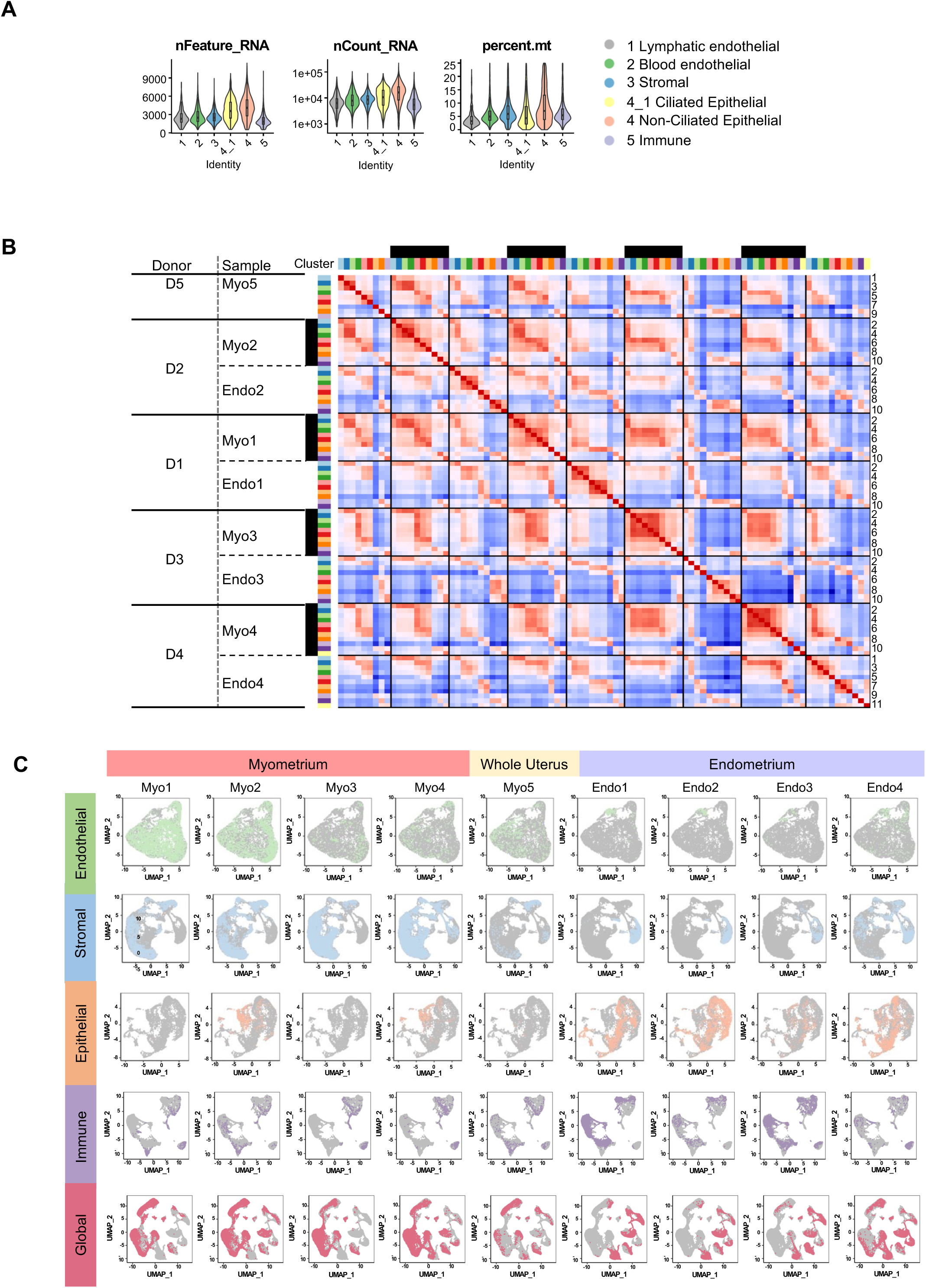
Quality control and evaluation of reproducibility of the nine datasets from the five healthy premenopausal donors. **A**. Distributions of the number of genes detected, library size, and percentage of mitochondrial reads across the major cell types, shown as violin plots, for past QC cells (<25% mitochondrial reads and >500 detected genes). **B**. Evaluation of cluster reproducibility across samples. Shown is the heatmap of pairwise centroid-centroid correlations among the 9-11 clusters obtained separately for each of the nine scRNA datasets. **C**. Comparison of the nine samples’ contributions (left to right) to the four major cell types (top four rows) and overall (bottom row). Shown in the UMAP plot for all the cells, with one cell type shown in each row (using a different color) and with the other cell types or in other samples shown in the background (color gray).

**Supplemental Figure 2.**
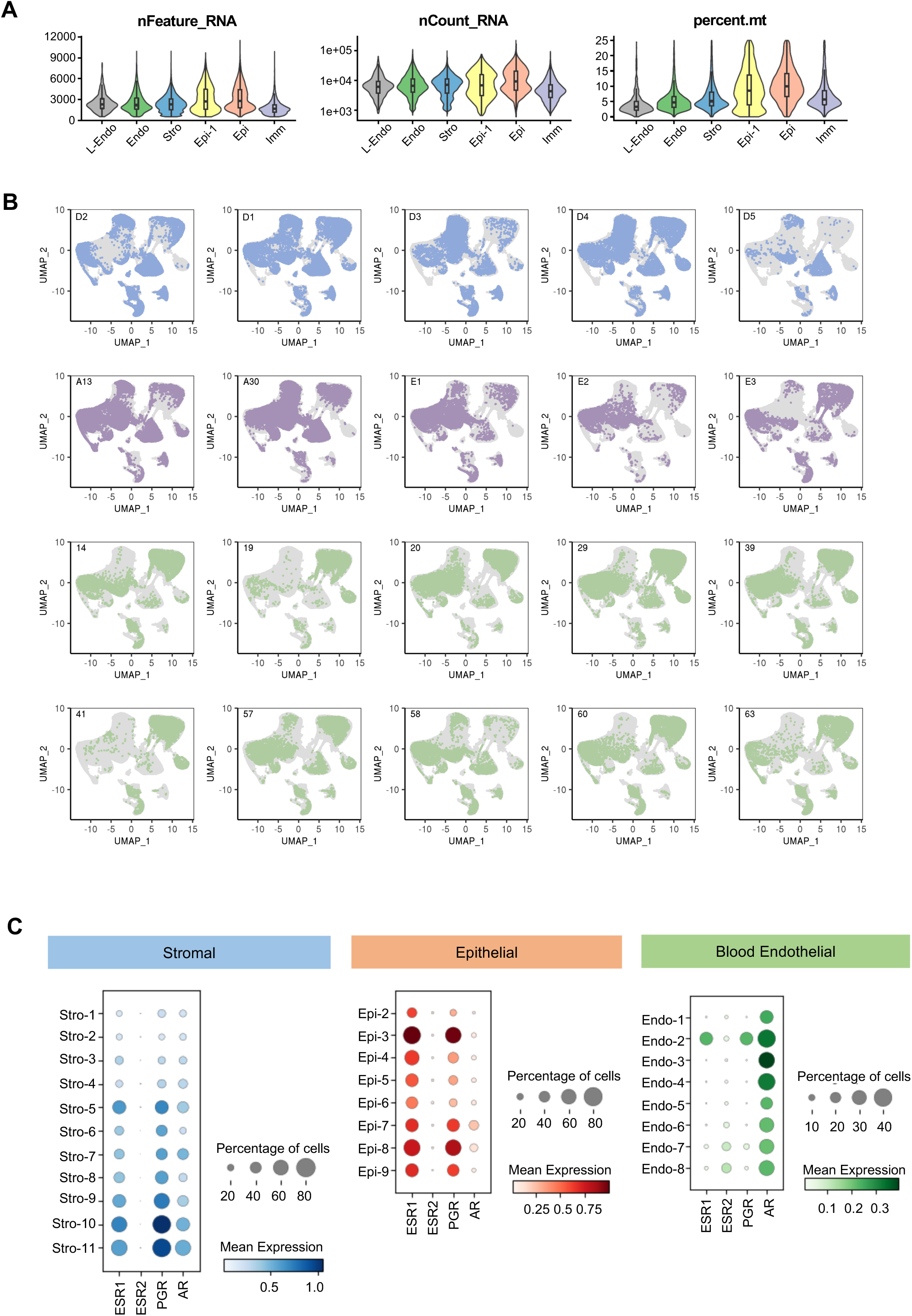
Quality control and evaluation of reproducibility of the 20 datasets; expression pattern of hormone receptors. **A**. Distributions of the number of genes detected, library size, and percentage of mitochondrial reads across the major cell types, shown as violin plots, for past QC cells. **B**. Comparison of the 20 samples’ contributions, shown in the same UMAP plot as in Figure 2A, with cells from each donor colored, and those from the other 19 samples shown in gray. **C**. Dot plots comparing the distribution patterns of hormone receptor genes (estrogen receptors ESR1. ESR2, progesterone receptor PGR, androgen receptor AR) across subtypes for the stromal, unciliated epithelial, and blood endothelial cells.

**Supplementary Figure 3.**
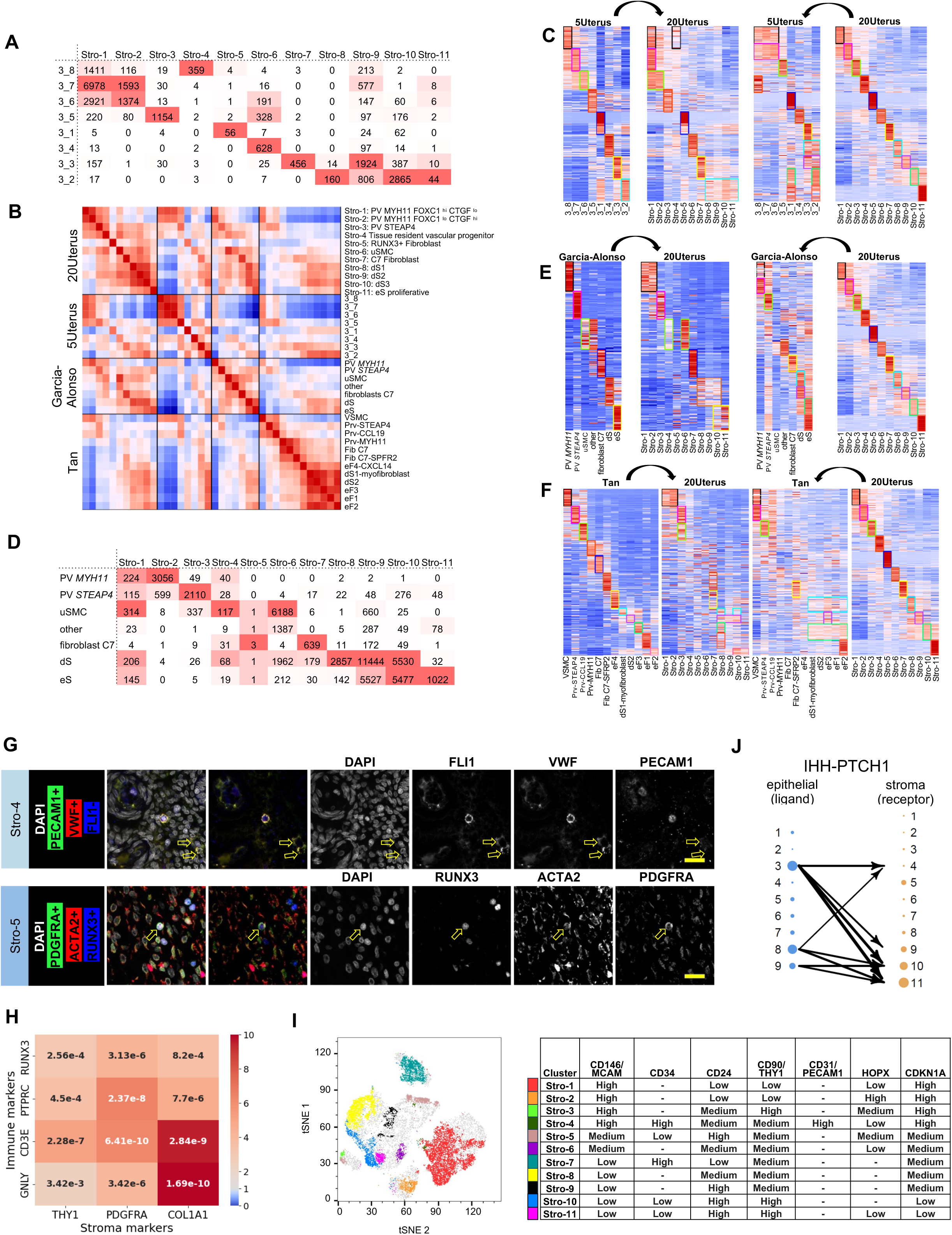
Multi-study integrated analysis and consensus annotation of 11 stromal cell subtypes. We compared stromal cell subclusters across four classification attempts: 8 subtypes in our 5-sample analysis, 11 subtypes in 20-sample joint analysis, 7 subtypes in the 15-sample analysis by Garcia-Alonso et al. (25), and 12 subtype centroids by Tan et al. **A**. Cross tabulation of stromal cell assignments between the 5-sample and 20-sample results, for cells in the 5 samples (also shown in Supplementary Table 4D). **B**. Centroid-centroid correlation across the four sets of stromal subtypes, shown as a heatmap of pairwise rank correlations (data included as Supplementary Table 8A). **C, E, F**. Subtype-specific differentially expressed genes obtained from one subtype set and compared across the centroids in one or more of the other classification results, to evaluate potential subtype matches. The pair of panels on the right are for 20-sample DE genes compared with the 5-sample, Garcia-Alonso et al. (25), and Tan et al. (26) subtypes. The pair of panels on the left of C-E are for DE genes from the 5-sample, Garcia-Alonso et al. (25), and Tan et al. (26) subtypes and compared with the 20-sampel centroids. All supporting data are in Supplemental Tables 8B-D. **D**. Cross tabulation of stromal cell assignments for cells in the 15 samples in Garcia-Alonso et al. (25) between the 20-sample results and the subtypes in Garcia-Alonso et al. (25) (also shown in Supplementary Table 4C). **G**. Immunofluorescence co-staining of a uterus sample using antibodies against unique markers for stromal subtypes. Upper panels: arrows indicate VWF+PECAM1+FLI1-cells, likely Stro-4. Lower panels: the yellow arrow indicates a RUNX3+ACTA2+PDGFRA+ cell, likely Stro-5. Scale bar: 20 um. **H**. Co-occurrence patterns between one of the immune markers (rows) and one of the stromal markers (columns) in Stro-5 cells, with the values shown as p-values for significant co-occurrence from Fisher’s Exact Test, and the color in the heatmap scaled with the odds ratio (red: OD>1, gray: OR=1). **I**. Seven-protein multiplexed measurements of ∼40,000 cells to validate the 11 stromal subtypes. The antibodies used are for MCAM, PECAM1, THY1, CD34, CD24, HOPX, and CDKN1A. Seven-channel signals were quantified by flow cytometry, with the tSNE projection performed in FlowJo v. 10.10. Eleven groups of manually defined cells (using criteria shown in the table) were visualized in the tSNE plot. **J**. Subtype-specific IHH signaling from epithelial cells to stromal cells. Shown is an “arrow plot” where high expression of the ligand IHH in one of the epithelial subtypes may be linked to high expression of the receptor PTCH1 in one of the stromal subtypes. Only the top 10% of the pairwise interactions were shown as arrows. Dot size for each subtype indicated its percent of cells expressing the ligand or the receptor.

**Supplemental Figure 4.**
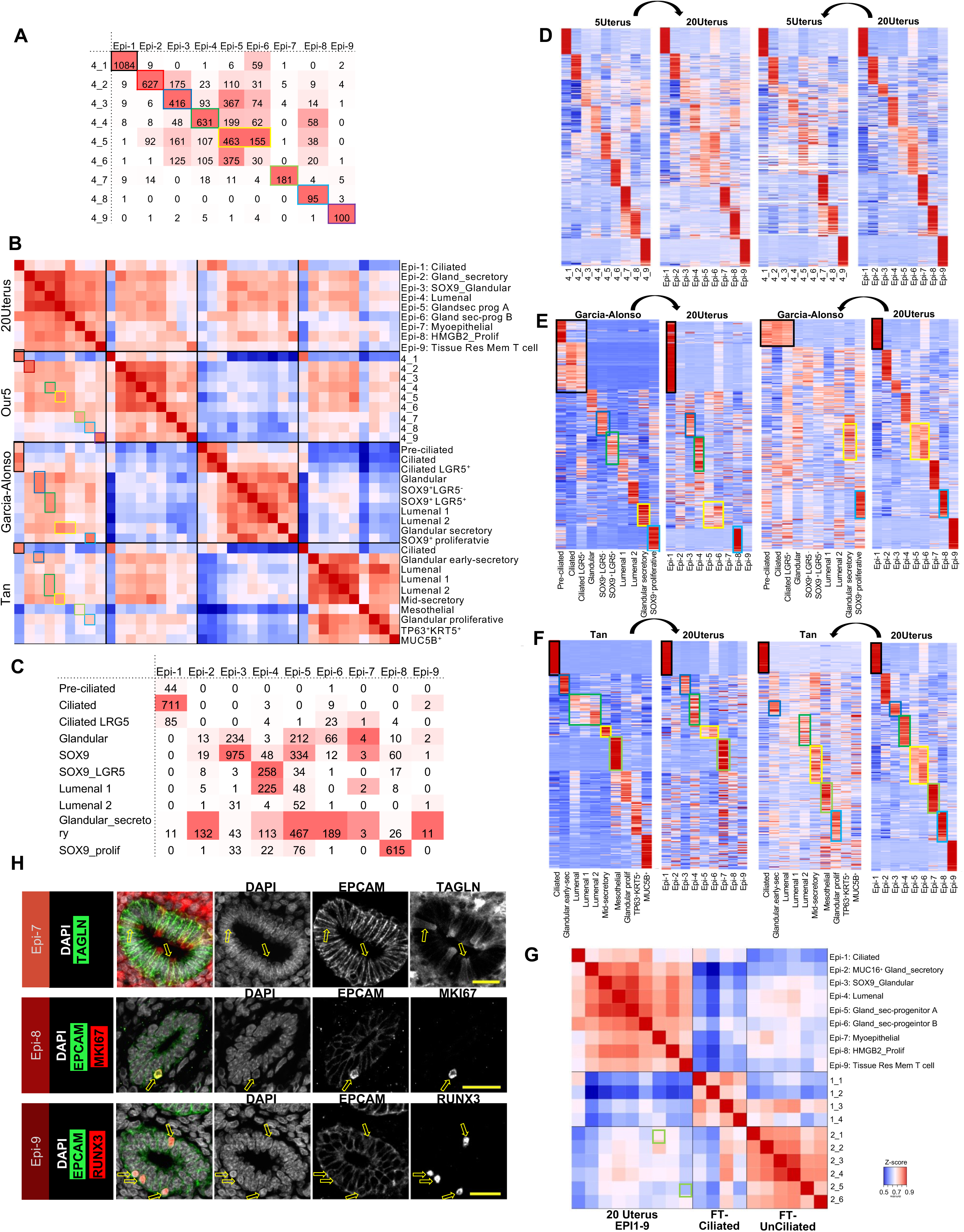
Integrated analysis and consensus annotation of 9 epithelial subtypes. We compared epithelial cell subtypes across four classification attempts: 9 subtypes in our 5-sample analysis, 9 subtypes in 20-sample joint analysis, 10 subtypes in the 15-sample analysis by Garcia-Alonso et al. (25), and 10 subtype centroids by Tan et al (26). **A**. Cross tabulation of epithelial cell assignments between the 5-sample and 20-sample results, for cells in the 5 samples (also shown in Supplementary Table 4D). **B**. Centroid-centroid correlation across the four sets of epithelial subtypes, shown as a heatmap of pairwise rank correlations (data included as Supplementary Table 9A). **C**. Cross tabulation of epithelial cell assignments for cells in the 15 samples in Garcia-Alonso et al. (25) between the 20-sample results and the subtypes in Garcia-Alonso et al. (25) (also shown in Supplementary Table 4C). **D-F**. Subtype-specific differentially expressed genes obtained from one subtype set and compared across the centroids in one or more of the other classification sets, to evaluate potential subtype matches. Method and format follow those described above for Supplemental Figure 3C, E, F, with data shown in Supplemental Tables 9B-D. **G**. Centroid-centroid comparisons between the 9 epithelial subtypes in uterus with the ciliated (1_1 to 1_4) and unciliated (2_1 to 2_6) cell subtypes reported previously for fallopian tube samples. Shown is the heatmap of rank correlation values. **H**. Immunofluorescence (IF) co-staining of a uterus endometrium sample using antibodies against unique markers for epithelial cell subtypes. Upper: TAGLN+ EPCAM+ cell, likely Epi-7. Middle: EPCAM+ MKI67+ cell, likely Epi-8. Lower: EPCAM+ RUNX3+ cells, likely Epi-9. Scale bar: 20 um.

**Supplemental Figure 5.**
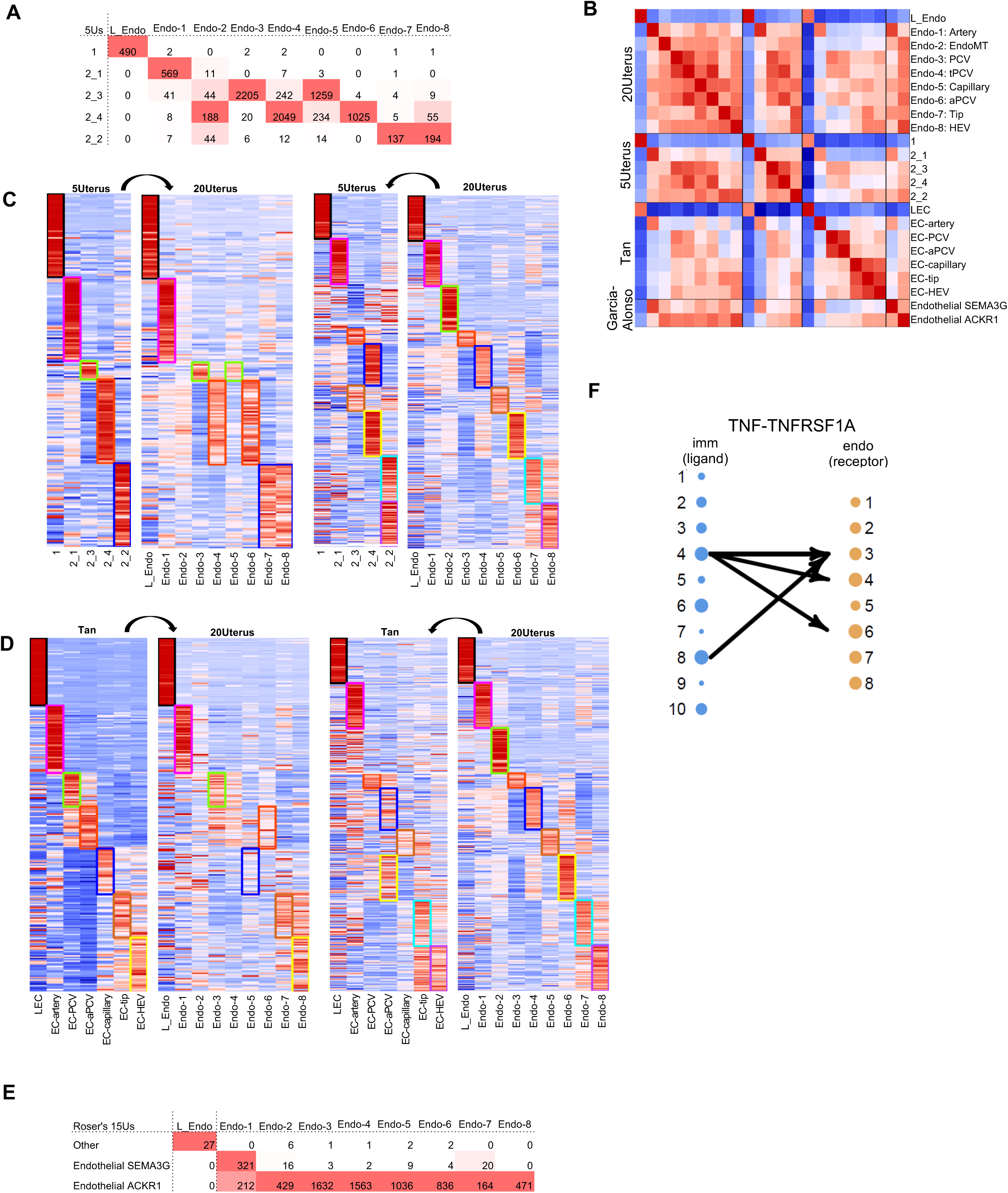
Integrated analysis and consensus annotation of eight blood endothelial subtypes. We compared endothelial cell subtypes across four classification attempts: 5 subtypes in our 5-sample analysis (4 are for blood endothelial), 9 subtypes in 20-sample joint analysis (8 for blood endothelial), two subtypes in the 15-sample analysis by Garcia-Alonso et al. (25), and seven subtype centroids by Tan et al (26). **A**. Cross tabulation of endothelial cell assignments between the 5-sample and 20-sample results, for cells in the 5 samples (also shown in Supplementary Table 4D). **B**. Centroid-centroid correlation across the four sets of endothelial subtypes, shown as a heatmap of pairwise rank correlations (data included as Supplementary Table 10A). **C-D**. Subtype-specific differentially expressed genes obtained from one subtype set and compared across the centroids in one or more of the other classification sets, for evaluating potential subtype matches. Method and format follow those described above for Supplemental Figure 3C, E, F, with data shown in Supplemental Tables 10B-C. We did not include a comparison with Garcia-Alonso et al. (25) as it had only two subtypes. **E**. Cross tabulation of endothelial cell assignments for cells in the 15 samples in Garcia-Alonso et al. (25) between the 20-sample results and the subtypes in Garcia-Alonso et al. (25) (also shown in Supplementary Table 4C). **F**. Subtype-specific TNF signaling from immune cells to endothelial cells. Shown is an “arrow plot” where high expression of the ligand TNF in one of the endothelial subtypes may be linked to high expression of the receptor TNFRSF1A in one of the stromal subtypes. Only the top 5% of the pairwise interactions were shown as arrows. Method and format are the same as described above in Supplemental Figure 3J.

**Supplemental Figure 6.**
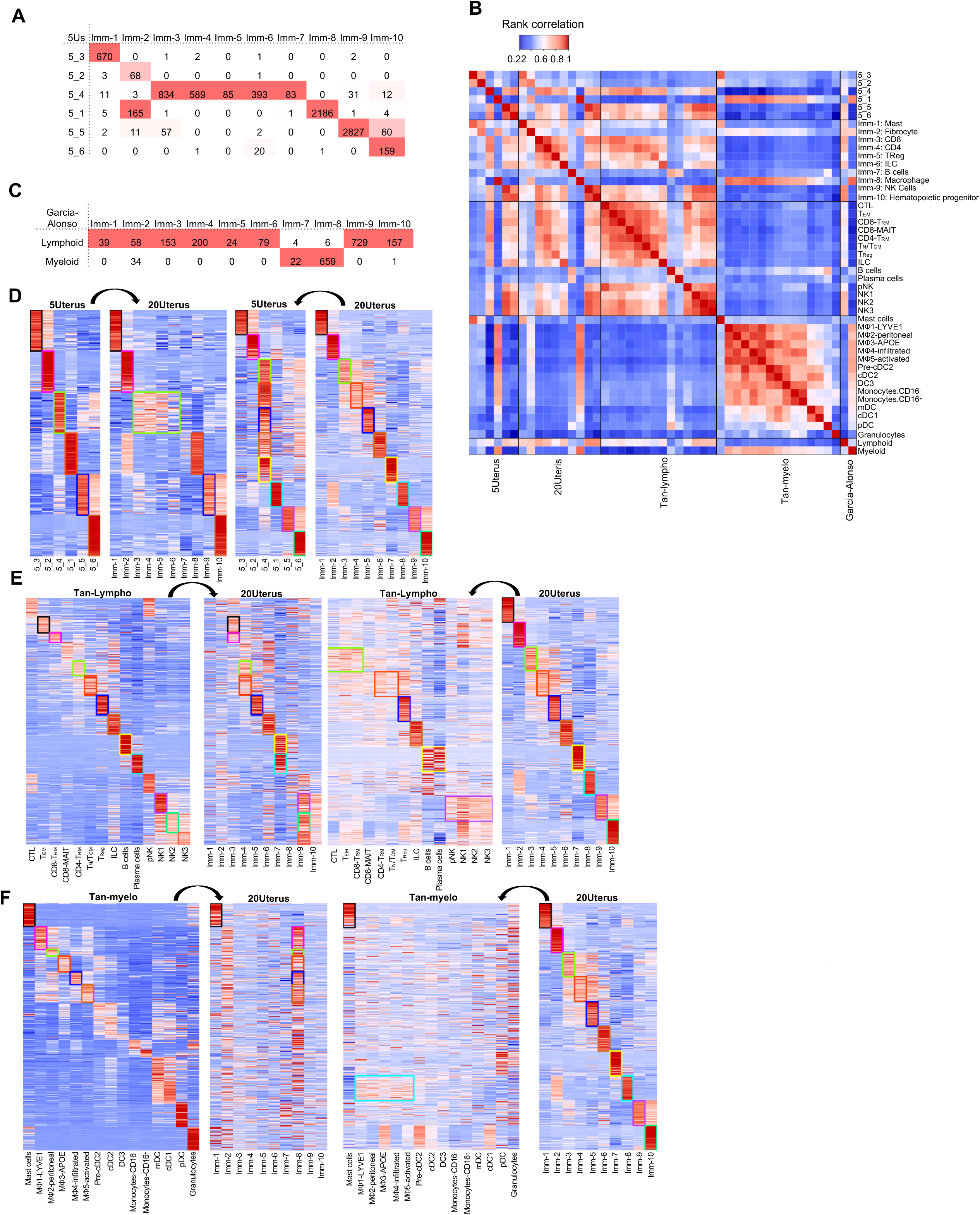
Integrated analysis and consensus annotation of 10 immune cell subtypes. We compared immune cell subtypes across five classification sets:6 subtypes in our 5-sample analysis, 10 subtypes in 20-sample joint analysis, 14 lymphoid subtypes and 15 myeloid subtypes in Tan et al. (26), and two subtypes in the 15-sample analysis by Garcia-Alonso et al. (25) **A**. Cross tabulation of immune cell assignments between the 5-sample and 20-sample results, for cells in the 5 samples (also shown in Supplementary Table 4D). **B**. Centroid-centroid correlation across the five sets of immune subtypes, shown as a heatmap of pairwise rank correlations (data included as Supplementary Table 11A). **C**. Cross tabulation of immune cell assignments for cells in the 15 samples in Garcia-Alonso et al. (25) between the 20-sample results and the subtypes in Garcia-Alonso et al. (25) (also shown in Supplementary Table 4C). **D-F**. Subtype-specific differentially expressed genes obtained from one subtype set and compared across the centroids in one or more of the other classification sets, for evaluating potential subtype matches. Method and format follow those described above for Supplemental Figure 3C, E, F, with data shown in Supplemental Tables 11B-D. We do not include a comparison with Garcia-Alonso et al. (25) since as it had only two subtypes.

## Supplemental Table Legends

**Table 1. Summary information of the five donors, uterus samples, and scRNA datasets.**

**1A. Metadata for the 11 uterus samples from the seven donors**. Included are donors’ source of collection, demographic information (Age, BMI, self-described race), menopausal status, and estimated cycling phase. From the seven donors, four had myometrium and endometrium samples analyzed separately, leading to 11 scRNAseq datasets. The post-QC cell counts and average number of detected genes per cell were included.

**1B. Cell counts for the five major cell types over the nine samples** from the five premenopausal donors, as assigned in the 20-sample joint analysis (see Methods).

**Table 2. Gene expression centroids from the five-sample analysis.**

**2A. Centroids for the five major cell types**. Shown are centroids values for the full set of 32,612 genes over the five major cell types, while splitting “4_1”, the Ciliated cells, from “4”, the non-ciliated epithelial cells. Values were the average of ln(CP10K+1) for cells in the cluster, where CP10K is counts-per-10K normalized values.

**2B. Centroids for the 28 subtypes.** Values are calculated in the same way as in **2A**, by the same gene order. Included are one subtype for lymphatic endothelial (not subclustered), and 4, 9, 8, and 6 subtypes for the blood endothelial, stromal, epithelial, and immune cells, respectively.

**Table 3. Markers for major cell types based on differential expression in the five-sample analysis.**

**3A. Centroid expression values of the top 150 differentially expressed (DE) genes** calculated for each of the major cell types (for ∼900 total), as shown in the heatmap in **Figure 1C**. The full set of DE genes was listed in 4B below, and ranked by fold-change when selecting the top 150. From the centroid data in 2A, the values for each gene were standardized by subtracting the mean and dividing by the standard deviation over the six centroid values.

**3B. List of computationally obtained DE genes for the major cell types**. Shown are p value, log2(Fold-change), percent detected in cells of the cluster (pct.1) and cells in all other clusters (pct.2), and the adjusted p value (p_val_adj), the indicated cell type, and the gene name. The number of genes vary by cluster due to FC>1.6 and other cutoffs (see Methods).

**3C-3F. List of DE genes for the subtypes** for the blood endothelial, stromal, non-ciliated epithelial, and immune subtypes, respectively. The content and format are the same as in 3B.

**Table 4: Metadata and cell counts by sample-subtype in the 20-sample joint analysis.**

**4A. Metadata for the 19 endometrium and 6 myometrium samples** (from 20 donors in 3 studies). Also included are cycle phase information and “Aggregated cycle” which combined some of the rare phases. To the right are cell counts over the five major cell types and 26 samples in the 20-sample joint clustering.

**4B. Cell count fractions by subtype, for each of the four major cell type and each of the 19 endometrium samples (ordered by phase)**. For each major cell type, the cell count fraction is calculated for each subtype, thus the total fraction for each major cell type is 100%.

**4C. Cross tabulation of cells’ assignment in the 15-sample analysis** between (1) the 39 subtypes in the 20-sample joint analysis and (2) the 21 cluster labels from Garcia-Alonso et al. (25). Note that the shifts across major cell types (in green) were minor when compared to those within major cell types (in red).

**4D. Cross tabulation of cells’ assignment in the 5-sample analysis** between (1) the 39 subtypes in the 20-sample joint analysis, same as in 4C, and (2) the 28 subtypes in the original 5-sample clustering.

**Table 5. Gene expression centroids from the 20-sample analysis.**

**5A. Centroids for the five major cell types**. Shown are centroids values for the full set of 43,090 genes, in the same format as in 3.

**5B. Centroids for the 39 subtypes.** Values are calculated in the same way as in **5A**, by the same gene order. Included are one subtype for lymphatic endothelial (not subclustered), and 8, 11, 9, and 10 subtypes for the blood endothelial, stromal, epithelial, and immune cells, respectively.

**Table 6. Markers for major cell types based on differential expression in the 20-sample analysis.**

**6A. List of computationally obtained DE genes for the major cell types. Format is the same as in** Table 3B. The number of genes vary by cluster due to FC>1.6 and other cutoffs (see Methods).

**6B-6E**. **List of DE genes for the subtypes** for the blood endothelial, stromal, non-ciliated epithelial, and immune subtypes, respectively. The content and format are the same as in 3B and 6A.

**Table 7: Gene-centroid values for literature-based markers** in subtypes for the four major cell types, as shown in **Figure 2C**. For each of the major cell types (endothelial, stromal, epithelial, and immune), a panel of marker genes was curated based on known cell type-specific function, prior knowledge of uterus tissue biology, and related pathologies. These genes were used, in conjunction with the centroids in Table 5B and DE genes in Table 6B-E, to annotate the identity of the cell subclusters. The values shown were based on centroids data in Table 5B and have been standardized for each gene (by subtracting the mean and dividing by the standard deviation).

**7A-7D**. Gene-Centroid values for literature-based genes over the endothelial, stromal, epithelial, and immune subtypes, respectively. They correspond to the four heatmaps shown in Figure 2C.

**Table 8. Stromal subtype annotation supporting data**, as shown in **Supplemental Figure 3B, C, E-F,** including centroid-centroid correlations and reciprocal marker expression comparison among the three studies,.

**8A. Pairwise rank correlation coefficients among subtypes, as shown in Supplemental Figure 3B**, for four sets of stromal cell classifications: 11 subtypes in the 20-sample analysis, 8 subtypes in the 5-sample analysis, 7 subtypes in Garcia-Alonso et al. (25), and 12 subtypes in Tan et al (26). The genes used for calculating correlations were obtained by (1) collecting top 50 DE genes for each stromal subtype in each of the four classifications, and (2) merging the DE genes into a union list.

**8B. DE genes selected from 20-sample results, compared between 20-sample centroids and three results**: 5-sample centroids, Garcia-Alonso centroids (25), and Tan et al(26). centroids. These correspond to the pairs of heatmaps on the right side of **Supplemental Figures 3C, 3E**, and **3F**, respectively. Top 50 DE genes with the largest fold-change were chosen for each of the 11 stromal clusters in the 20-sample analysis, and plotted across the four sets of centroid results. When fewer than 50 DE genes pass the threshold, all of them are used. Values have been standardized for each gene. Marker selection and standardization were applied in the same way for 8C-8E below.

**8C. DE genes selected from 5-sample results, compared between 5-sample centroids and the 20-sample centroids**, corresponding to the pair of heatmaps on the left side of **Supplemental Figures 3C.**

**8D. DE genes selected from Garcia-Alonso et al. (25) results, compared between Garcia-Alonso et al. (25) centroids and the 20-sample centroids**, corresponding to the pair of heatmaps on the left side of **Supplemental Figures 3E.**

**8E. DE genes selected from Tan et al. (26) results, compared between Tan et al. (26) centroids and the 20-sample centroids**, corresponding to the pair of heatmaps on the left side of **Supplemental Figures 3F.**

**Table 9. Epithelial subtype annotation supporting data**, including centroid-centroid correlations and reciprocal marker expression comparison among the three studies, as shown in **Supplemental Figure 4B, D-F**.

**9A. Pairwise rank correlation coefficients among subtypes, as shown in Supplemental Figure 4B**, for four sets of epithelial cell classifications: 9 subtypes in the 20-sample analysis, 9 subtypes in the 5-sample analysis, 10 subtypes in Garcia-Alonso et al. (25), and 10 subtypes in Tan et al. (26). The genes used for calculating correlations were obtained in the same way as in 8A: by (1) collecting top 50 DE genes for each epithelial subtype in each of the four classifications, and (2) merging the DE genes into a union list.

**9B. DE genes selected from 20-sample results, compared between 20-sample centroids and three results**: 5-sample centroids, Garcia-Alonso (25) centroids, and Tan et al. (26) centroids. These correspond to the pairs of heatmaps on the right side of **Supplemental Figures 4D, 4E**, and **4F**, respectively. Top 50 DE genes with the largest fold-change were chosen for each of the 9 epithelial clusters in the 20-sample analysis, and plotted across the four sets of centroid results. Methods and format are the same as in Table 8B-E. Values have been standardized for each gene.

**9C. DE genes selected from 5-sample results, compared between 5-sample centroids and the 20-sample centroids**, corresponding to the pair of heatmaps on the left side of **Supplemental Figures 4D.**

**9D. DE genes selected from Garcia-Alonso et al. (25) results, compared between Garcia-Alonso et al. (25) centroids and the 20-sample centroids**, corresponding to the pair of heatmaps on the left side of **Supplemental Figures 4E.**

**9E. DE genes selected from Tan et al. (26) results, compared between Tan et al. (26) centroids and the 20-sample centroids**, corresponding to the pair of heatmaps on the left side of **Supplemental Figures 4F.**

**Table 10. Endothelial subtype annotation supporting data**, including centroid-centroid correlations and reciprocal marker expression comparison among the three studies, as shown in **Supplemental Figure 5B-D**.

**10A. Pairwise rank correlation coefficients among subtypes, as shown in Supplemental Figure 5B**, for four sets of endothelial cell classifications: 9 subtypes in the 20-sample analysis, 5 subtypes in the 5-sample analysis, 2 subtypes in Garcia-Alonso et al. (25), and 7 subtypes in Tan et al. (26). The genes used for calculating correlations were obtained in the same way as in 8A: by (1) collecting top 50 DE genes for each endothelial subtype in each of the four classifications, and (2) merging the DE genes into a union list.

**10B. DE genes selected from 20-sample results, compared between 20-sample centroids and two other results**: 5-sample centroids and Tan et al. (26) centroids. These correspond to the pairs of heatmaps on the right side of **Supplemental Figures 5C** and **5D**, respectively. Methods and format are the same as in Table 8B-E. Values have been standardized for each gene.

**10C. DE genes selected from 5-sample results, compared between 5-sample centroids and the 20-sample centroids**, corresponding to the pair of heatmaps on the left side of **Supplemental Figures 5C.**

**10D. DE genes selected from Tan et al. (26) results, compared between Tan et al. (26) centroids and the 20-sample centroids**, corresponding to the pair of heatmaps on the left side of **Supplemental Figures 5D.**

**Table 11. Immune subtype annotation supporting data**, including centroid-centroid correlations and reciprocal marker expression comparison among the three studies, as shown in **Supplemental Figure 6B, D-F**.

**11A. Pairwise rank correlation coefficients among subtypes, as shown in Supplemental Figure 6B**, for five sets of immune cell classifications: 10 subtypes in the 20-sample analysis, 6 subtypes in the 5-sample analysis, 14 lymphoid and 15 myeloid subtypes in Tan et al. (26), and 2 subtypes in Garcia-Alonso et al (25). The genes used for calculating correlations were obtained in the same way as in 8A: by (1) collecting top 50 DE genes for each immune subtype in each of the five classifications, and (2) merging the DE genes into a union list.

**11B. DE genes selected from 20-sample results, compared between 20-sample centroids and three results**: 5-sample centroids, 14 lymphoid, and 15 myeloid subtypes centroids in Tan et al. (26). These correspond to the pairs of heatmaps on the right side of **Supplemental Figures 6D, 6E**, and **6F**, respectively. Methods and format are the same as in Table 8B-E. Values have been standardized for each gene.

**11C. DE genes selected from 5-sample results, compared between 5-sample centroids and the 20-sample centroids**, corresponding to the pair of heatmaps on the left side of **Supplemental Figures 6D.**

**11D. DE genes selected from 14 lymphoid subtypes in Tan et al. (26), compared between Tan et al. (26) centroids and the 20-sample centroids**, corresponding to the pair of heatmaps on the left side of **Supplemental Figures 6E.**

**11E. DE genes selected from 15 myeloid subtypes in Tan et al. (26) compared between Tan et al.** (26) **centroids and the 20-sample centroids**, corresponding to the pair of heatmaps on the left side of **Supplemental Figures 6F.**

## Supplemental Methods

### Experimental Model and Subject details

#### Healthy human specimen collection

Full thickness uterus samples were obtained from five premenopausal, one postpartum, and one peri-menopausal women (ages 28-52; Supplemental Table 1A). Four of the subjects were undergoing surgery for benign indications, and samples were obtained from the University of Michigan Reproductive Subject Registry and Sample Repository (RSRSR)(95). The collection of tissue was approved by the University of Michigan Institutional Review Board (HUM00167998). The remaining three subjects were from organ donors without overt diseases in the reproductive system, procured by the International Institute for the Advancement of Medicine (IIAM) through standard procedures. As the tissues from these deceased donors were de-identified, their use does not fall under regulated human subject’s research. In accordance with their standard procedures, the donors were prepped for organ recovery while maintained on a ventilator. During the procedure the sternum was opened to enable access to the thoracic and abdominal cavities which were packed with ice. The ventilator was turned off prior to aortic cross-clamp and an arterial line was opened to start the flow of one of three preservation media: Belzer UW®Cold Storage Solution (Bridge of Life, SC, USA), Custodiol® HTK (Histidine-Tryptophan-Ketoglutarate) Solution (Essential Pharmaceuticals, NC, USA), or SPS-1 Static Preservation Solution (Organ Recovery Systems, IL, US)). This serves to flush the organs and maintain their viability during recovery. The uterus was surgically removed and delivered to the lab within 24 hours. Based on the information provided by IIAM these donors had 46XX karyotypes, negative infection and pregnancy screens, and were without known disease in the reproductive system such as endometriosis, significant uterine surgery, systemic disease or previous obstetric complications. The menstrual cycle phase for all samples collected in this study were determined by a gynecologic pathologist based on histologic evaluation and blood work analysis.

#### Ethical approval process

The IIAM procures tissue and organs for non-clinical research from Organ Procurement Organizations (OPOs), which comply with state Uniform Anatomical Gift Acts, are certified and regulated by the Centers for Medicare and Medicaid Services. These OPOs are members of the Organ Procurement and Transplantation Network and the United Network for Organ Sharing (UNOS) and operate under a set of standards established by the Association of Organ Procurement Organizations and UNOS. Written informed assent from the next-of-kin of the deceased donor was obtained. A biomaterial transfer agreement is in place between IIAM and the authors that restricts the use of the tissue for pre-clinical research. The use of deceased donor tissue in this research is categorized as ‘not regulated’, per 45 CFR 46.102 and the ‘Common Rule’, as it does not involve human subjects and complies with the University of Michigan’s IRB requirements (HUM00250736).

### Method Details

#### Tissue processing and preparation for single-cell RNA sequencing

Surgical samples were collected in Hank’s Balanced Salt Solution (HBSS) in the operating room. The samples were transported at room temperature to the lab and processed immediately for dissociation. The cadaveric samples were processed within 24-36 hours of removal due to logistical constraints, including travel and availability of core laboratory services (see Supplemental Table 1). If a delay in processing for dissociation was needed, the sample was maintained at 4°C in tissue preservation media (UW or HTK). For most samples, the uterus was dissected into two anatomic segments: endometrium and myometrium.

For the endometrium samples, the tissue was incubated in 5 mM EDTA in HBSS (depleted of calcium and magnesium) on ice for 30 minutes. The tissue is then filtered through a 100-micron strainer. For each 100-200mg of tissue, samples were resuspended in 18mg of pronase (Sigma P5147) diluted in 10ml of Opti-mem (Gibco 31985-062) and incubated at <u>37°C for 5 minutes</u>. The pronase treated tissue samples were filtered using a 100 micron strainer, and tissue chunks were transferred to 10ml digestion buffer 2 (containing <u>1 mg/ml</u> collagenase D (Roche 11088866001), 1 mg/ml hyaluronidase (Sigma H3884), and 2 U/ml DNase I in HBSS), and incubated for 15 minutes at 37°C. The supernatant from the digested cells was collected and quenched with ∼3ml of 10% fetal bovine serum (FBS). The remaining undigested tissue was subjected to a second round of digest. Dissociated cells from the two rounds of digest were washed 2x in 10% FBS/DMEM, centrifuged at 600g for 5 minutes, and resuspended in 1ml 10% FBS/DMEM and left on ice while preparing myometrial samples.

For myometrium samples, 100-200mg of tissue were resuspended in 18mg of pronase (Sigma P5147) diluted in 10ml of Opti-mem (Gibco 31985-062) and incubated at <u>37°C for 10 minutes</u>. The pronase treated tissue samples were filtered through a 100-micron strainer, and tissue chunks were transferred to 10ml digestion buffer 2 (containing <u>1.5 mg</u>/ml collagenase D (Roche 11088866001), 1 mg/ml hyaluronidase (Sigma H3884), and 2 U/ml DNase I in HBSS) and incubated for 30 minutes at 37°C. The supernatant was collected, stored on ice and quenched with 3ml of 10% FBS/DMEM. While the undigested tissue fragments were subjected to 3x rounds of digest (incubation times for the last 3 rounds of digest were 30, 20 and 20 minutes, respectively). After 4 rounds of sequential digest, the quenched supernatant from each sample was pooled and washed in PBS. Cell suspensions were resuspended in EasySep RBC depletion reagent (Stemcell Technologies #18170) or Red Blood Cell Lysis Solution (Miltenyi Biotec # 130-094-183) according to their manufacturer’s instruction. After centrifugation at 600g for 5 minutes, cells were washed with 0.04% BSA/PBS 2-3 times and filtered through a 100 μm filter. Samples were resuspended in 1 ml 10% FBS/DMEM and live cells were collected by flow cytometry, and submitted to the University of Michigan Advanced Genomics Core for processing on the 10X Genomics Chromium platform. Sequencing libraries were sequenced by Illumina NovaSeq to create 151-nt reads for the transcript.

#### FFPE Immunofluorescence

Paraffin sections from the uterus were obtained from the University of Michigan Reproductive Subject Registry and Sample Repository (See Supplemental Table 2 for patient information – Note patient samples include sequenced and non-sequenced individuals). Five-micron FFPE tissue sections were deparaffinized in Histoclear 3x for 5 minutes, followed by serial EtOH washes from 100% to 30%, 3 minutes each. After the 30% ethanol, the slides were transferred into two containers of deionized water for 3 min each. Tissue sections were subsequently permeabilized in 0.1% Triton in PBS for 15 minutes. For all antibodies, antigen retrieval was performed by boiling in 10mM sodium citrate, pH 6.0 for 30 minutes. Sections were blocked in 1xPBS supplemented with 3% BSA and 500mM glycine for two hours at room temperature. Endogenous peroxidases and alkaline phosphatases were blocked by a ten-minute incubation in BloxAll solution (VectorLabs, Cat. No: SP-6000). The primary antibodies used include PECAM1, EPCAM-488 (Abcam, ab237395, 1:200), RUNX3 (Sigma HPA059006, 1:100), smooth muscle actin (ACTA2-594, Abcam ab202368, 1:200), PSMB2 (Abcam, ab137108, 1:100), VWF (Abcam, ab11713, 1:100), FLI1(Abcam, ab133485, 1:100), and PDGFRA (R&D Systems, AF1062-SP, 1:50). All secondary antibodies were conjugated to either Alexa-488-, Alexa- 555-, or Alexa-647 (LifeTechnologies/ Molecular Probes) and diluted to1:1000. DAPI was used as a nuclear counterstain. The images were taken with a Nikon A1R-HD25 confocal microscope and processed with ImageJ. Staining was generally done in a combination as indicated in each panel.

#### Flow Cytometry Analysis

Endometrial and myometrial samples were dissociated as described above and resuspended in phosphate buffered saline (BD Biosciences 223142). 1-5 million single-cells were stained with an Fc blocking antibody (<u>BD Pharmingen™ Human BD Fc Block™</u> Catalog #: 564219). After blocking cells were stained with a multiplex antibody combination including both primary antibodies and/or isotype controls. The primary and isotype control antibodies are listed in the table below:

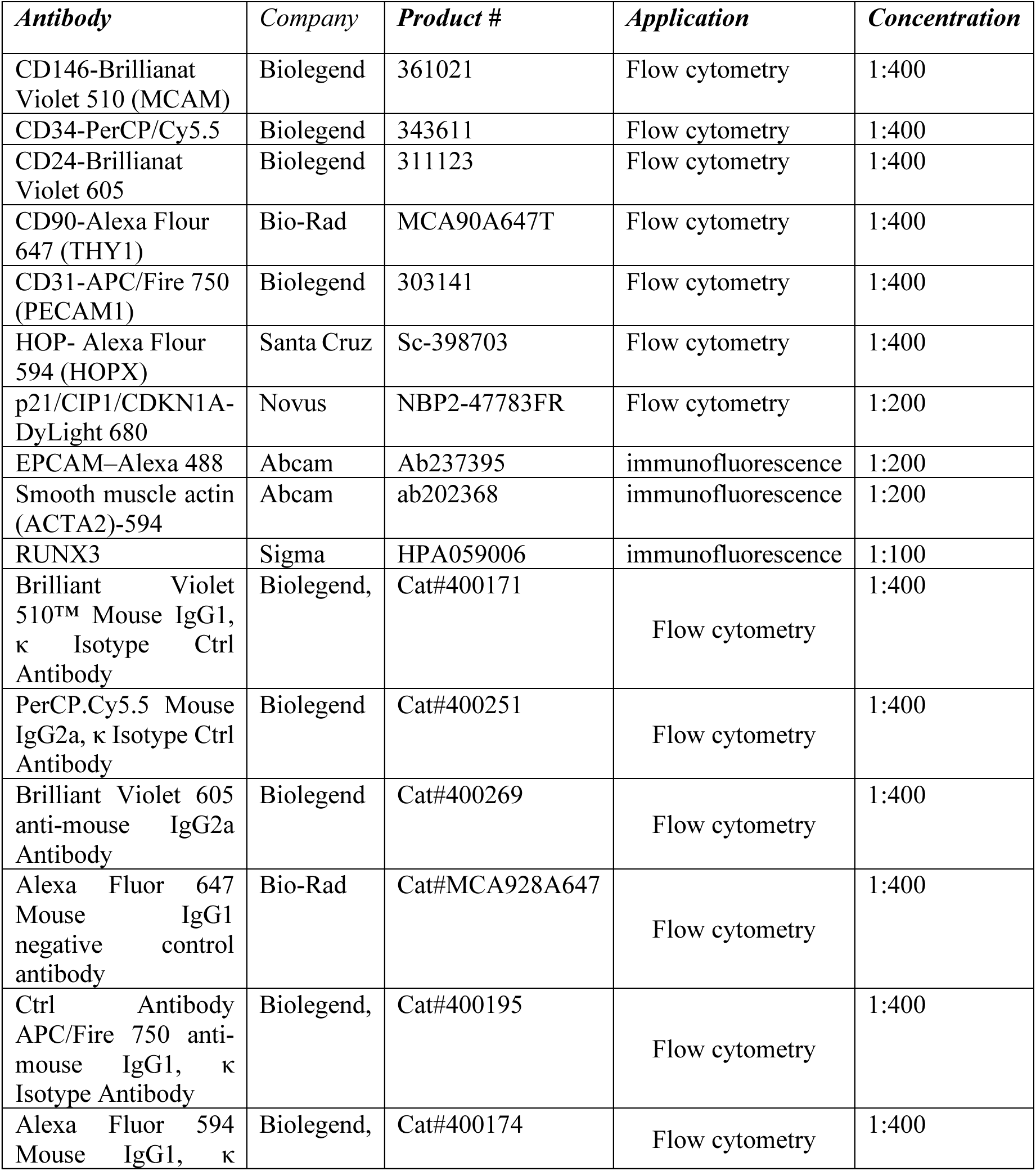
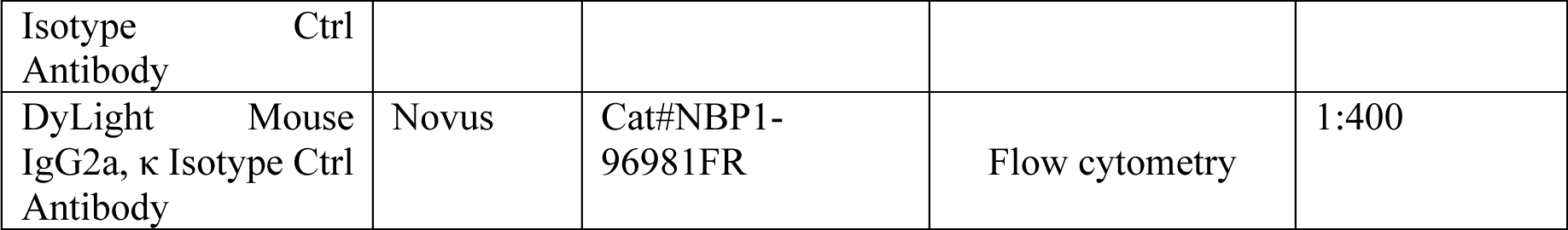

A seven multiplex flow cytometry requires compensation and the precise delineation of positive and negative gate thresholds, but this is challenging because of spectral overlap between fluorochromes and the complexity of biological samples. To overcome these challenges, we ran "Flow Minus One" (FMO) controls to determine the lower threshold for positive gates and the upper threshold for the negative gates for each of the seven antibodies. For each FMO control, 200-300k cells are stained with 6/7 fluorochromes used in the experiment except for the antibody of interest. Therefore, for the 7 antibodies we had 7 FMO controls. Additionally, we had three replicates for the pooled antibody sample. Using both FMOs and the multiplexed antibody cocktail sample, we determined the gates for the positive and negative signals. Since flow cytometry measures the fluorescent intensity for each of the 7 proteins (negative, low medium and high) on millions of cells, we performed unbiased clustering on the millions of analyzed cells to identify the different stromal cell subtypes of cells. To determine whether the 11 stromal subpopulations, we identified by single cell RNAseq map can be identified in the flow data, we pseudo colored the cells with the following expression patterns below (see table below).

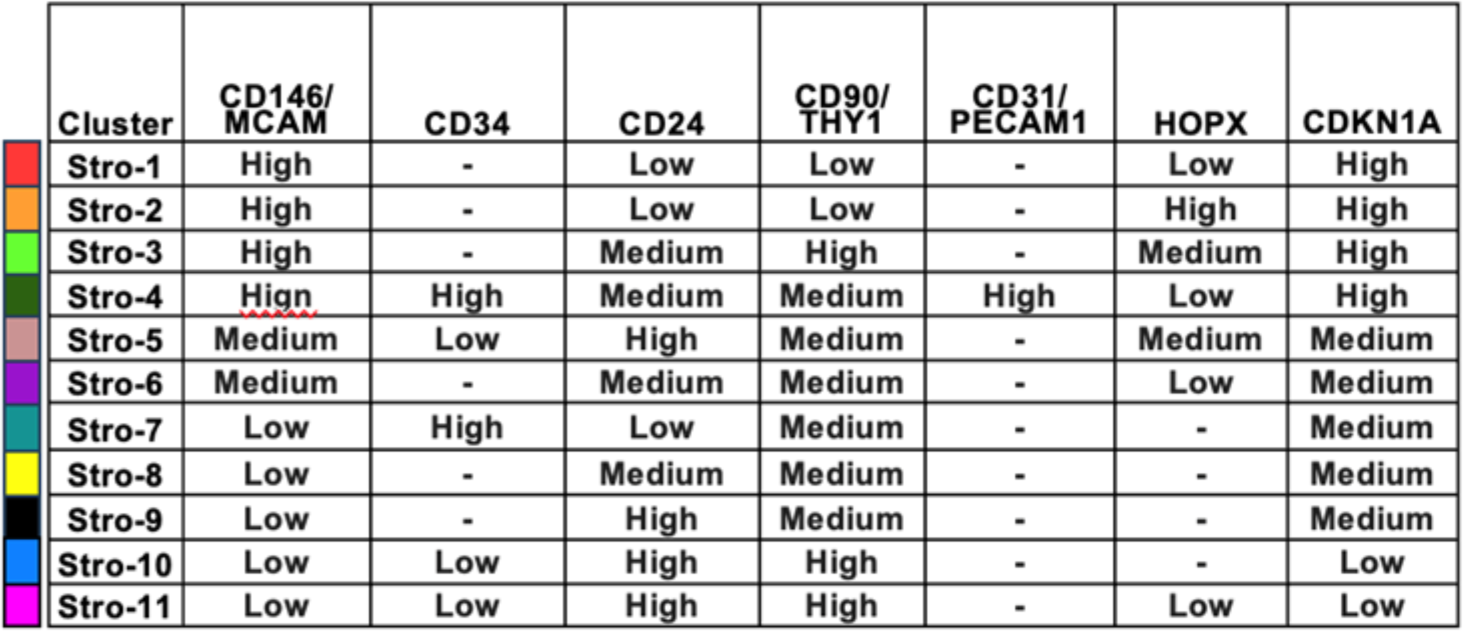

Using these expression patterns for the seven markers – we resolved all 11 subtypes in the unbiased flow cytometry defined clusters.

### Sequencing data availability

The raw sequence data are deposited at GEO under accession # GSE260658, with reviewer token uvkhqmioltmrrop.

### Computational methods

#### Single-cell RNA-seq data processing

We produced 11 scRNAseq datasets from the seven human subjects (Supplemental Table 1A). Four of the five healthy premenopausal donors produced two datasets each, for endometrium and myometrium. For the fifth healthy donor and the postpartum donor, the endometrium was extremely thin; and we analyzed one full-uterus sample consisted of mostly myometrium cells. The healthy postmenopausal subject produced one myometrium dataset.

The 11 tissue samples were prepared with the 10x Chromium Controller and sequenced on an Illumina NovaSeq 6000. The reads were aligned to human reference genome GRCh38, with the cell barcodes and UMIs processed by Cell Ranger v4.0.0 (or newer), which generated cell-by-gene count matrices for each sample.

Cells with fewer than 500 detected genes or with >= 25% of detected transcripts corresponding to mitochondrial-encoded genes were filtered out. In the nine datasets from the five healthy premenopausal donors, 1,810 - 8,161 cells passed QC, with 23,327 - 29,648 genes detected over all cells in each sample. For the postpartum donor, 5,161 cells passed QC, with 24,474 genes detected overall, and an average of 2,171 detected genes and 7,716 UMIs per cell. For the postmenopausal donor, 5,085 cells passed QC, with 26,259 genes detected overall, and an average of 2,330 detected genes and 7,813 UMIs per cell. See Supplementary Table 1A.

#### Recurrently applied data processing steps

We mostly used *Seurat* (V4.1.1;(96)) for processing the scRNAseq data.

For each cell, raw UMI counts are log-normalized using the *Seurat* function NormalizeData, which (1) divides by the total number of UMIs per cell, (2) multiplies by 10,000 to obtain a transcripts-per-10K (CP10K) measure, and (3) log-transforms by ln(CP10K+1).

For projection and clustering, the normalized data were standardized by gene using the *Seurat* function ScaleData, which centers by the mean and scales by standard deviation, (E-mean(E))/sd(E), with a ceiling of 10 imposed (to mitigate the effects of small standard deviation values). We will specify when non-standardized data are used.

The selection of highly variable genes (HVGs) was done before gene standardization, using the *Seurat* function FindVariableFeatures with default parameters. This step models the relationship between gene mean and variance using LOESS, uses this to perform an idealized “standardization” of the genes, then selects the top 2000 HVGs. For PCA, UMAP, and t-SNE projections we use default parameters in *Seurat*, while manually choosing the number of PCs for use in UMAP and t-SNE projections and in clustering.

#### Evaluation of between-sample and between-subject reproducibility

For the nine datasets from the five healthy premenopausal donors, we standardized the gene expression matrix, selected 2,000 HVGs to perform PCA, selected the top 9-14 PCs based on the elbow of the scree plot, and computed t-SNE and UMAP projections. We then performed Louvain-Jaccard clustering using *Seurat* function FindClusters, varying the "resolution" parameter from 0.1 to 1 in steps of 0.1 to obtain 9 - 11 clusters for each dataset. We calculated pairwise rank correlation coefficients between all cluster centroids from all datasets (Supplemental Figure 1B).

#### Integrating data over multiple samples

For the five healthy donors, the endometrium and myometrium data of the same donor were directly merged, as they were processed in the same experiment. To account for batch effects among donors, we use the CCA-based data integration pipeline in *Seurat,* using donor ID as the batch factor. This involves two *Seurat* functions, FindIntegrationAnchors and IntegrateData. In FindIntegrationAnchors we use k.filter = min(n,200), and in IntegrateData we use k.weight = min(n,100), where n represents the smallest number of cells from any donor involved in the integration. Otherwise we employ default parameters.

#### Identification of major cell types for five healthy premenopausal uterus donors

We obtained integrated data for the five-uterus dataset, containing a total of 50,992 cells that pass QC, with an average of 2,755 detected genes and 10,626 UMIs per cell. A total of 32,612 genes were detected in this integrated dataset.

We next standardized the dataset, performed PCA using 2,000 HVGs, and used the top 14 PCs for tSNE and UMAP projections and Louvain-Jaccard clustering. We initially detected 15 clusters and, by using marker genes, we annotated cluster 1 as lymphatic endothelium, cluster 2 as blood endothelium, clusters 3-8 as stromal, clusters 9-11 as epithelium, and clusters 12-15 as immune. The cell counts are: C1 lymphatic endothelial - 513, C2 blood endothelial - 8,678, C3 stromal - 26,694, C4 epithelial - 6,465, and C5 immune - 8,653. Figure 1B shows the UMAP projection with these clusters labeled. Focused UMAPs of each major cell type appear in Figure 1E, and the contributions from the nine datasets are in Supplemental Figure 1C.

#### Focused re-clustering to discover cell subtypes

For cells in clusters C2 - C5 we performed re-clustering to discover subtypes within each major cell type. C1 contained too few cells (513) and we did not re-cluster them. The recurring workflow is to extract all normalized cells for the cluster, select 2,000 HVGs, standardize the genes, perform PCA, determine top PCs by the scree plot, compute tSNE and UMAP projections, and run Louvain-Jaccard clustering using resolutions from 0.1 to 1 (in steps of 0.1). We used resolution of 0.2 for C3 and 0.1 for the others. This results in four C2 subclusters (using the top five PCs), eight C3 subclusters (using the top eight PCs), six C4 subclusters (using the top six PCs), and six C5 subclusters (using the top eight PCs).

Based on markers, one of the six C4 clusters (4_1) corresponded to ciliated cells (1,168 cells), while the other five are unciliated epithelium (5,286 cells). One of the five is large: 3,236 (61% of the unciliated cells); the other four contain an average of 513 cells each. Therefore, we further extracted and subclustered the full unciliated subset following the outline above. This resulted in 8 unciliated epithelial clusters. UMAP projections for the four major cell types are in Figure 1E.

#### Integrated analyses of 20-uterus donors by combining scRNAseq data from Garcia-Alonso et al. (25) 2021 and Wang et al. 2020 (19)

We combined our 5-uterus donors with the 15 samples analyzed in Garcia-Alonso et al. 2021 (25). Five of the 15 were first reported in Garcia-Alonso et al.(25) while the other 10 were first presented in Wang et al. 2020 (19).

Data for the five donors by Garcia-Alonso et al. 2021 (25) came as six samples (one had endometrium and myometrium analyzed separately), in 11 raw gene expression matrices (2-3 for each). They were downloaded from E-MTAB-10287 (https://www.ebi.ac.uk/biostudies/arrayexpress/studies/E-MTAB-10287). We filtered cells by the same QC process, removing those with <500 detected genes or >= 25% of detected transcripts corresponding to mitochondrial-encoded genes. After CP10K normalization and log transformation, we merged samples from the same donor to obtain five datasets. There are 52,341 cells that pass QC, with 27,699 genes detected overall, and an average of 2,313 detected genes and 8,557 UMIs per cell.

For the 10 10x samples in Wang et al. 2020(19), we download the cell-by-gene count matrix and corresponding annotations from GSE111976 (https://ftp.ncbi.nlm.nih.gov/geo/series/GSE111nnn/GSE111976). After the same QC and normalization steps there are 64,577 cells, with 29,666 genes detected, and an average of 2,632 detected genes and 10,153 UMIs per cell. All ten were endometrial samples.

The cell type annotations for the 15 samples by Garcia-Alonso et al. (25) were downloaded from https://www.reproductivecellatlas.org/non-pregnant-uterus.html. Due to differences in pre-processing steps, not all pass-QC cells in our analyses were assigned cell type labels in Garcia-Alonso et al. (25)

We put the pass-QC and normalized data from the 20 samples into one *SeuratObject*, selected 2,000 HVGs, and corrected for batch using the integration pipeline in *Seurat,* with donor ID as the batch factor. The resulting integrated dataset contains 167,910 cells and 43,090 detected genes, with an average of 2,570 detected genes and 9,799 UMIs per cell. Nineteen of the donors contributed endometrial samples and six contributed myometrial samples. The menstrual cycle phase for each donor is also included as part of the dataset; we aggregate the provided phase information into five categories (proliferative, early secretory, early-mid secretory, mid secretory, and late secretory) in Figure 2C. This and other metadata are in Supplemental Table 4.

#### Identification of major cell types in the 20-uterus dataset

We standardized the integrated data, performed PCA, and chose the top 14 PCs to compute tSNE and UMAP projections (Figure 2A, Supplementary Figure 2B). With resolution 0.3, Louvain-Jaccard clustering produced 17 initial clusters. Like before, we used known markers to annotate clusters 1-4 as epithelial, clusters 5-12 as stromal, cluster 13 as blood endothelial, cluster 14 as lymphatic endothelial, and clusters 15 - 17 as immune cells. The last cluster, cluster 16, apparently contains immune, stromal, and epithelial cells, and it was examined further by its own round of re-clustering. This proceeded in the following manner: extract the log-normalized expression data for all cells in cluster 16 from donors that contribute more than 30 cells (this eliminates one out of the 20 donors), integrate the remaining 19 samples, standardize the integrated gene values, perform PCA on the integrated data, restrict to the eight top PCs, compute tSNE and UMAP projections, and run Louvain-Jaccard clustering. A 12-subcluster result separated the epithelial cells as 16_5, and stromal cells as 16_4. We thus have the epithelial cells in clusters 1-4 and 16_5, stromal cells in clusters 5-12 and 16_4, and immune cells in clusters 15, 17, and the remainder of 16 after removing 16_4 and 16_5. The cell numbers for major cell types (C1 lymphatic endothelial - 548 cells, C2 blood endothelial - 16,648 cells, C3 stromal - 89,281 cells, C4 epithelial - 45,715 cells, C5 immune - 15,718 cells) and subtypes are in Supplemental Table 4. Figure 2A contains a UMAP projection of all 167K cells with these major cell types labeled.

#### Focused re-clustering to identify cell subtypes in 20-uterus dataset

To re-cluster cells within a major cell type, we removed any donors that contributed fewer than 30 cells to the cluster, then integrate the log-normalized expression data from the remaining samples in the subset. Next perform PCA on the integrated data, manually select top PCs, and find tSNE and UMAP projections.

For re-clustering C2 blood endothelial cells, 28 cells from two donors from Wang et al. (19) were removed. Two of the subclusters, Endo-1 and Endo-2, were stable across different Leiden subclustering parameters and were set aside (as "done"). The remaining blood endothelial were preprocessed by standardizing the genes and computing new PCA, tSNE, and UMAP projections. Leiden clustering led to seven clusters, one of which (501 cells) was subsequently removed due to lower library sizes and higher mitochondrial content. The final, eight-cluster solution, Endo-1 though Endo-8, contains 16,119 blood endothelial cells.

For C3 stromal cells, initial Louvain-Jaccard clustering revealed 10 subclusters, one of which (3,406 cells) was removed due to low quality of the cells. After adding cluster 16_4 (a total of 85,875 cells), we obtained a new set of 10 clusters, one of which was noted for high expression of RUNX3 (an immune cell marker). This cluster, when examined alone, split into two populations: one with RUNX3 expression (now called Stro-5) and the other with no expression (now called Stro-4). In all we have 11 stromal cell subtypes, Stro-1 through Stro-11.

For C4 epithelial cells, Donor D5 contributed only 24 cells, which were removed. Clustering the remaining 45,691 cells showed six clusters. One of these, Epi-1 (4,475 cells), showed clear correspondence with the ciliated cells. For the unciliated epithelial cells, we noted that the 10 samples from Wang et al. (19) provided 31,455 cells, much more than the 5,440 cells from our five samples and the 4,321 cells from the five donors in Garcia-Alonso et al. (25). We decided to down-sample all epithelial cells (34,397 cells) from Wang et al. (19) by first obtaining a provisional eight-cluster solution. One of these eight clusters consisted of ciliated cells, so we kept up to 2,000 cells from each of the other seven clusters, resulting in 10,884 unciliated cells (down from 30,804). When combined with those from our five samples and the five from Garcia-Alonso et al. (25), we have 20,645 unciliated epithelial cells.

Initial re-clustering of the down-sampled unciliated population led to six clusters, two of which were stable, and set aside. The remaining 18,420 unciliated cells were further clustered into 6 clusters, leading to Epi-2 through Epi-9 (Epi-1 contains the ciliated cells).

For C5 immune cells, an initial clustering produced seven subclusters, three of which (later called Imm-8, Imm-9, and Imm-10) were well separated in PCA and were set aside, with the remaining cells further subclustered into 12 subclusters. One of these 12 clusters (1062 cells) was removed due to low library size and high mitochondrial content, and the rest were merged into seven clusters, leading to the final set of 10 clusters (Imm-1 through Imm-10) containing 14,656 cells.

Figure 2B shows UMAP plots for C2-C5, with subclusters labeled by color. Figure 2D shows the subcluster composition of each major cell type, further stratified by the stage in the menstrual cycle and the origin (endometrium or myometrium) of the sample (Supplemental Table 4B contains the cell type composition information for each donor). The table below summarizes the initial and final cell counts.

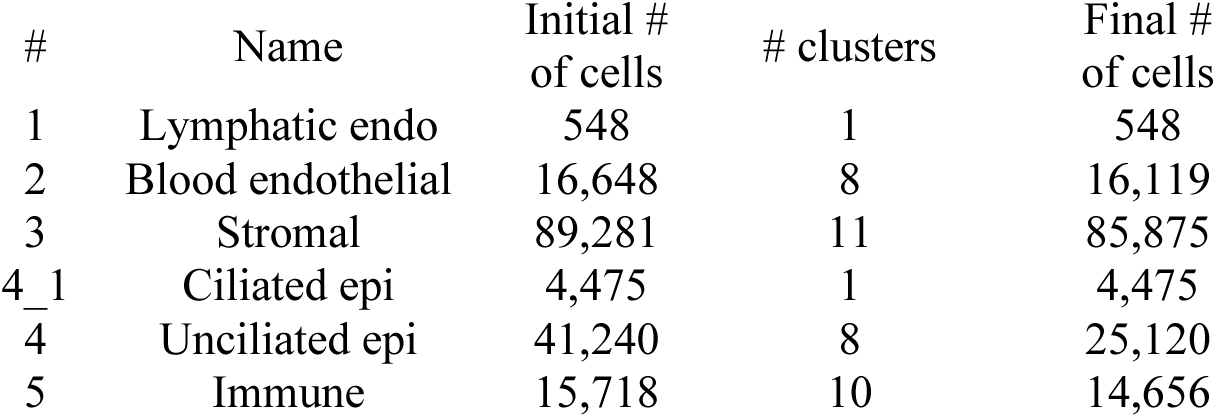

#### Computation of centroids

Centroids of clusters (or subclusters) were computed using the *Seurat* function AverageExpression, with default parameters, followed by adding one and taking the natural log. Briefly, the CP10K values were averaged over the cells in the cluster, then we take ln(avg + 1) as the centroid. Centroids for the 5-sample and the 20-sample datasets were in Table 2 and Table 5, respectively.

For the 15-samples in Garcia-Alonso et al.(25) we also calculated centroids to produce the centroid-centroid correlations shown in Supplemental Figures 3B, 4B, 5B, and 6B. Instead of using CP10K, we started from the 15-uterus integrated gene expression data produced in that study (downloaded from https://www.reproductivecellatlas.org/non-pregnant-uterus.html) and used cell type assignments from that study. Similarly, centroids for Tan et al. (26) were computed using the gene expression h5ad objects from that study (downloaded from https://singlecell.jax.org/datasets/endometriosis-2022). For example, the Tan et al. (26) stromal centroids used in Supplementary Figure 3 were computed using the file endo-2022_stromal.h5ad. For Supplemental Figure 4G, centroids for the subtypes of epithelial cells in the fallopian tubes were provided by the authors (51).

#### Differentially expressed genes (DEGs)

DEGs for a given cluster were obtained by comparing cells in that cluster against cells in all others clusters, using the Wilcoxon Rank Sum test implemented by the *Seurat* function FindAllMarkers. The criteria are (1) the gene must be detected in at least 10% of cells in the cluster of interest, and (2) there must be at least a 1.6-fold change. DEGs for cell subtypes were calculated in reference to other subtypes that belong to the same major cell type. Results were included in Table 3 and Table 6.

In several cases, we employ a stronger definition of DEGs than that presented above. In these figures, we restrict to all DEGs that have a fold change larger than 2 (ave_log2FC >= 1) and that have a difference of at least 20% in detection rate inside the target cluster vs outside (that is, pct.1 – pct.2 >= 0.2, using the column names from Table 6A). This definition is used in Figure 1C (where the 150 markers with the largest fold change are included for each of the major cell types) and in the supervised assignment of the D6 (postpartum) and D7 (postmenopausal) cells to the major cell types from the 20 uterus integrated dataset (further information below).

DEG lists for the cell types in Garcia-Alonso et al. 2021 (25) were found in Supplemental Table 6A from that work. DEG lists for the cell types in Tan et al. 2022(26) were downloaded from https://singlecell.jax.org/datasets/endometriosis-2022. For Supplemental Figure 4G, DEGs for the fallopian tube are obtained from Supplemental Tables 4A and 5A of Ulrich et al., 2022 (51). For Figure 1C, we chose 150 DEGs with the largest fold change.

The expression patterns of DEGs in the centroids of subclusters are visualized in heatmaps in Figures S3C,E,F, S4D-F, S5C,D, and S6D-F. In these panels, the 50 markers with the greatest fold change for each subcluster are selected for visualization. If any of the selected markers are not detected in one of the datasets, these markers are not included in the heatmaps.

#### Using RNA Velocity analysis to infer differentiation relationships

For the 50,992 cells in the five-sample dataset we obtained the spliced and unspliced transcript counts by using velocyto.py (v0.17.17) and using the GRCh38 genome annotation and repeat mask annotation files. The resulting .loom file was analyzed in python using *scVelo* (V0.2.6;(97)). For Figure 3, the RNA velocity came from cells in the 5-sample data, and the PCA projections came from the 20-sample data (we replace the projections computed in scVelo with our own projections). As t-SNE and UMAP often fail to maintain long-range relationships (for more "distant" cell types) we only present PCA for *scVelo* visualization.

#### Cell annotation of the postmenopausal and postpartum samples

Cells in the postpartum (D6) and postmenopausal (D7) samples were assigned to major cell types using the healthy sample centroids (Table 6A) based on the highest rank correlation, which were calculated for each cell using the stricter set of DEGs. In addition to fold change >= 2 and at least 20% difference in detection rate we further selected 100 DEGs with the largest fold change (600 genes for 5 major types plus Epi-1). Of these, 581 were detected in D6 and 593 in D7, and they were used as features for the supervised assignment. The Epithelial population was assigned no cell from D6 and only 15 from D7 (7 ciliated, 8 unciliated), thus we did not summarize their phase-pattern in Figure 4B.

The assignments of D6 and D7 cells to the subtypes were carried out in a similar fashion, using HVGs within the major cell type rather than DEGs. The cell type compositions of D6 and D7 are presented in Figure 4.

#### Curation and analyses of relevant makers from literature

We curated panels of known markers to aid in cell annotation (Table 7) as they complement DEGs from computational analysis (Tables 3 and 6). They were collected by searching relevant literature for both disease pathology and cell function. The expression patterns of some of the most notable literature-based markers were visualized in the major cell types of the five uterus dataset in a dot plot in Figure 1D and in the subclusters of the 20 uterus dataset as a heatmaps in Figure 2C. Finally, expression patterns of several hormone receptors (ESR1, ESR2, PGR, and AR) were compared across cell subtypes in dot plots in Supplemental Figure 2C.

#### Feature selection for centroid-centroid rank correlations

We used pairwise rank correlation to examine the similarities among cluster centroids (Figures 3G, S1B, S3B, S4B, S4G, S5B, and S6B). Most of these used the 50 DEGs with the largest fold change. For example, for Figure S3B (for stromal subclusters from four datasets), we obtained the top 50 DEGs for the five-uterus stromal subclusters (Table 3D), the 20-uterus subclusters (Table 6C), the Garcia-Alonso et al. (25) subclusters that we deemed “stromal,” and the stromal subclusters from Tan et al. 2022(26). The union of these gene lists were used for the final correlation calculation in Figure S3B.

#### Ligand-receptor analysis

We consider the list of ligand-receptor pairs identified in Ramilowski et al. 2015 (98) and downloaded from https://github.com/LewisLabUCSD/Ligand-Receptor-Pairs/blob/master/Human/Human-2015-Ramilowski-LR-pairs.txt. We restrict to our analysis to the 2,006 pairs ligand - receptor pairs detected in the 20-uterus dataset.

For a fixed ligand-receptor pair, we quantify the communication from cluster A to cluster B using an interaction score that is obtained by multiplying the expression of the ligand in centroid of A by the expression of the receptor in the centroid of B. We use log-normalized centroids, as provided in Supplementary Tables 3 and 6.

To visualize the interactions scores for one ligand-receptor pair between a specific set of clusters, we create an “arrow plot,” as in Supplemental Figures 3J and 5F. To do this, the clusters of interest that express the ligand are represented as dots on the left side of the plot, and the clusters expressing the receptor are represented as dots on the right side of the figure. The size of the dot is related to the percent of cells in the cluster that express the gene in question. We draw an arrow from a dot on the left to a dot on the right to represent the interaction score between the two clusters: the weight of the arrow is proportional to the interaction score. In Figures S3J and S5F, only the interactions in the top 10% and 5% (respectively) of interaction scores are represented with arrows.

Figure 3H is a summary of all interaction scores. There is a heatmap for each pair of major cell types. For each of these heatmaps, we restrict to the top 5% of all relevant interaction scores, compute the sum of the remaining interaction scores for every subcluster combination, and standardize the resulting values.

#### Co-expression analysis

For Stro-5 cells, we sought to evaluate if they co-express both immune and stromal markers, focusing on 3-4 canonical markers for each category. Since the transcript counts are sparse, many of these markers are not detected in many of the Stro-5 cells. We therefore turned the task into an analysis of present-absent patterns, i.e., rather than calculating the correlation of expression levels between Gene-1 and Gene-2 over all Str-5 cells, we scored each cell as having these genes to be either present (detected with at least one reads) or absent (not detected), and asked if Gene-1 and Gene-2 are both detected in Stro-5 cells more than by chance. This is quantified by using a 2-by-2 contingency table, where the expected rate of both-Present is the product of the rate of Present for the two genes. This led to a p-value from the Fischer’s Exact Test for each gene pair, and an odds ratio (OR) where OR>1 means the two genes tend to co-occur, while OR<1 means they tend to be mutually exclusive, reflecting a scenario when some cells express stromal markers while other cells express immune markers. Supplemental Figure 3H summaries the evidence of co-occurrence between three stromal makers and four immune markers, with p values shown, and with colors indicating OR, where all ORs are greater than 1

